# Distinct forms of dopamine transmission control locomotion and learning

**DOI:** 10.64898/2026.07.24.740623

**Authors:** Matthew M McGregor, Andrew G. Yee, Selin Ekici, Saige K. Power, Riccardo Melani, Joy Adler, Christina S. Winborn, Matthew J. Kennedy, Nicolas X. Tritsch, Christopher P. Ford

## Abstract

The neuromodulator dopamine is essential for voluntary movement and learning from experience. While these behaviors often occur simultaneously and rely on overlapping nigrostriatal dopamine circuits, they are also separable and can manifest independently. How a single neuromodulatory system controls such disparate aspects of behavior in parallel remains poorly understood. Here we show that striatal dopamine is released through two spatially and functionally distinct modes that are independently regulated and serve distinct behavioral roles. Eliminating dopamine release underlying bulk extracellular accumulation reveals a spatially restricted mode of transmission that maintains phasic dopamine receptor activation on striatal projection neurons. We find that selective loss of diffuse dopamine impairs striatal circuit excitability and reduces locomotion, while point-to-point transmission is sufficient to maintain striatal spine density and support motor and associative learning. We propose that the geometry of dopamine release determines its behavioral output, a principle that explains the breadth of dopamine’s functions and more broadly elucidates how neurons multiplex different forms of information.

## Main Text

Nigrostriatal dopamine neurons control both voluntary movement and learning by regulating striatal output. Dopamine has been implicated in the initiation and invigoration of movement^1–6^, the acquisition and consolidation of motor sequences^7–13^, and the moment-to-moment modulation of striatal circuit gain^14–17^. The expansive axonal arborization of dopamine neurons within the striatum^18^, and the low density of release sites^19,20^ and synaptic specializations^21–23^ have long been taken as evidence that dopamine signals through volume transmission by escaping the perisynaptic space to produce gradual changes in extracellular concentrations across large volumes of tissue. How a spatially indiscriminate signal supports this behavioral diversity is currently unresolved. One possibility is that molecular heterogeneity across dopamine neuron subtypes accounts for this diversity through selective targeting of downstream populations^24,25^. However, overlapping striatal innervation patterns^24^ and the demonstration that neighboring axons can encode opposing behavioral variables^3^ suggest that rapid decoding mechanisms that operate over sub-cellular spatial scales may also contribute. Thus, despite dopamine’s established behavioral specificity, how striatal circuits decode different dopamine signals remains unclear.

Recent evidence suggests that in addition to volume transmission, dopamine neurons can also engage in spatially restricted, point-to-point transmission, where high local concentrations of dopamine phasically activate receptors at defined postsynaptic targets^26–28^. Both the spatial dynamics of release and rapid onset of dopamine receptor activation of point-to-point transmission appear inconsistent with the slowly evolving, low-concentration extracellular dopamine transients produced by spillover^28–30^. Despite this evidence, the assumption has been that the source of dopamine underlying both modes is similar in regulation and downstream effects, differing only in spatiotemporal profile. This however has been difficult to evaluate, as the approaches frequently used when examining striatal dopamine measure release dynamics but not the activation of postsynaptic receptors. Whether the sources and regulation of dopamine signals contributing to these different modes are in fact independent, and whether their divergent spatiotemporal properties enable the differential encoding of distinct behaviors, has yet to be addressed.

We sought to address these two possibilities using a combination of approaches to measure both the presence and postsynaptic encoding of dopamine. Using conditional and viral knockouts of the active zone scaffold proteins RIM1 and RIM2, which localize voltage-gated calcium channels near release machinery to drive high-probability dopamine release^31–35^, we found that it was possible to selectively eliminate spatially diffuse release while preserving point-to-point transmission. We found that loss of spatially diffuse release produced alterations in striatal spiny projection neuron (SPN) excitability and reduced gross locomotion, while point-to-point transmission, which remained intact, was associated with preserved dendritic spine density and normal motor learning. These findings reveal independently regulated forms of dopamine transmission and suggest that dopamine neurons multiplex information through the geometry of release to simultaneously regulate multiple aspects of behavior.

### Spatially diffuse release is not required for phasic activation of dopamine receptors

The canonical view of dopamine signaling predicts that perturbations that reduce spatially diffuse release should produce proportional reductions in postsynaptic receptor activation (Fig. 1A). To test this, we perturbed dopamine release with a battery of manipulations. For each, we independently measured dopamine spillover in the extracellular space using fast-scan cyclic voltammetry (FSCV) and point-to-point transmission to D2-receptor (D2R) expressing indirect pathway SPNs (iSPNs). D2R activation was measured by virally expressing exogenous G protein-coupled inwardly rectifying potassium (GIRK) channels, which provide a rapid, analog, membrane-delimited, and Gβγ-mediated readout of receptor signaling in the form of inhibitory post-synaptic currents (D2-IPSCs)^28–30^ (Fig. 1A). These D2-IPSCs are dependent upon high, local concentrations of dopamine^28^. If spatially diffuse release drives receptor activation, then both measures should co-vary proportionally across manipulations (Fig. 1A).

**Figure 1:**
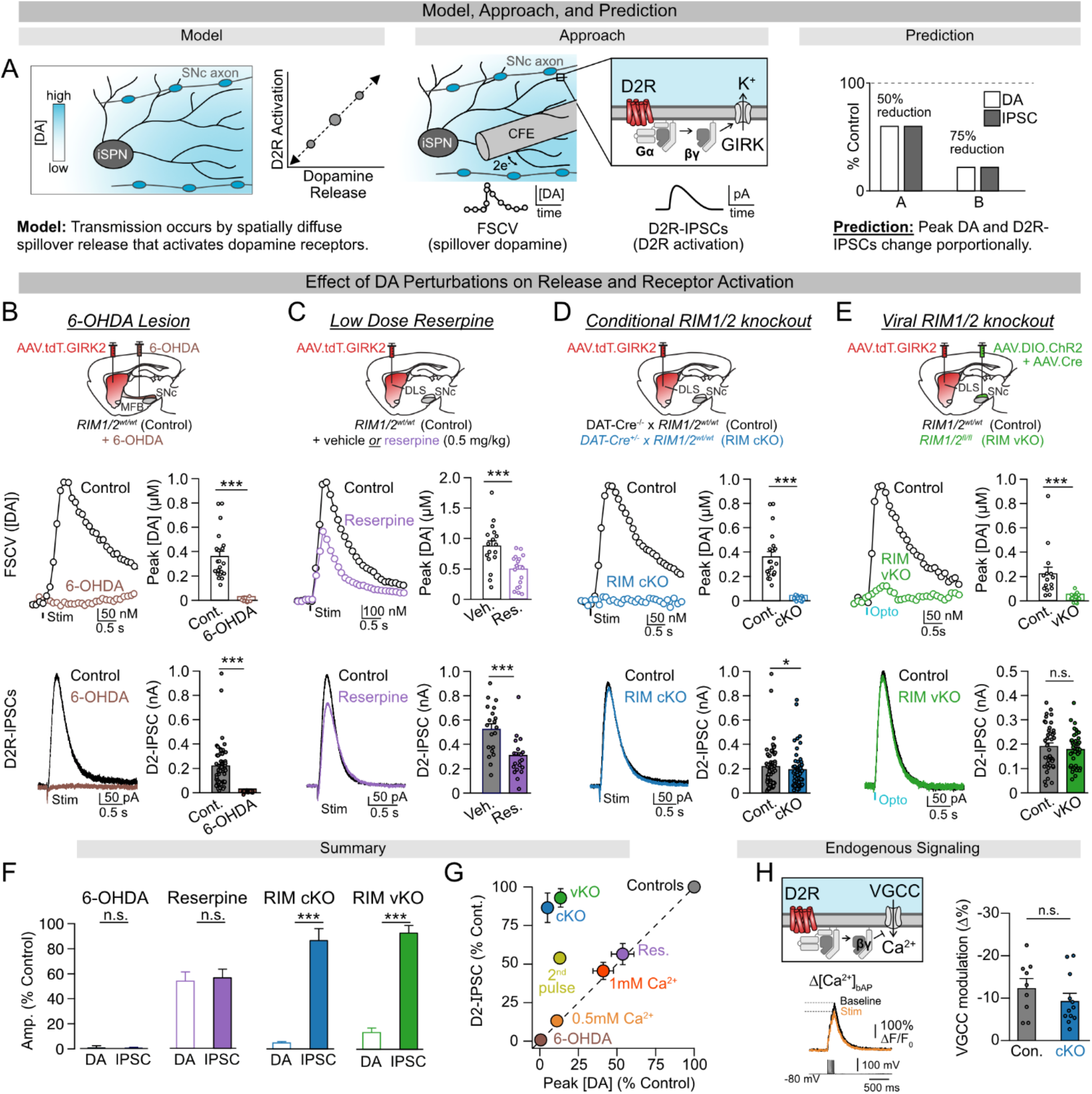
Spatially diffuse dopamine release is not required for dopamine receptor activation. A. Schematic depicting the canonical model of volume transmission by substantia nigra (SNc) dopamine neurons, methodological approach to test it, and prediction of how dopamine receptor activation changes in relation to spatially diffuse release. B. Effect of lesioning dopamine neurons (6-OHDA) on the amplitude of dopamine transients (*** p<0.001) and D2R-IPSCs (***p<0.001). Wilcoxon rank-sum test. C. Effect of low dose reserpine administration (0.5mg/kg, Res.) on the amplitude of dopamine transients (*** p<0.001) and D2R-IPSCs (***p<0.001). Wilcoxon rank-sum test. D. Effect of conditionally knocking out RIM1/2 in dopamine neurons (RIM cKO, cKO) on the amplitude of dopamine transients (*** p<0.001) and D2R-IPSCs (*p<0.05). Wilcoxon rank-sum test. E. Effect of virally knocking out RIM1/2 in dopamine neurons of adult animals (RIM vKO, vKO) on the amplitude of dopamine transients (*** p<0.001) and D2R-IPSCs (n.s. p>0.05). Wilcoxon rank-sum test. F. Summary of relative effects of dopaminergic lesioning (n.s. p>0.05), low-dose reserpine administration (n.s. p>.05), conditional RIM1/2 knockout (*** p<0.001), and viral RIM1/2 knockout (*** p<0.001) on the amplitude of dopamine transients and D2R-IPSCs, Wilcoxon rank-sum test. G. Summary of the relative effects of dopaminergic lesioning, altering extracellular calcium, low-dose reserpine administration, conditional and viral RIM1/2 knockout, and short-term plasticity on the amplitude of dopamine transients and D2-IPSCs. H. Schematic of D2R modulating voltage gated calcium channels (VGCC) through βγ-mediated signaling (top left). Example of calcium influx in iSPNs evoked by backpropagating action potentials (bAP) and the reduction following electrically stimulated dopamine release (bottom left). Summary of VGCC modulation in control (Con.) and RIM1/2 knockout mice (cKO, n.s. p>0.05). Wilcoxon rank-sum test. Summary data are the mean ± SEM.

We began with perturbations expected to produce broad reductions in dopamine release. Lesioning nigrostriatal inputs with 6-OHDA (Fig 1B, F), partially reducing vesicular content with a low dose of reserpine (0.5 mg/kg) (Fig. 1C, F), and lowering release probability with stepwise reductions in extracellular calcium (Suppl. Fig S1) all reduced both evoked extracellular dopamine transients and D2-IPSCs to a similar extent. The proportional reduction in extracellular dopamine and receptor activation (Fig. 1G) is thus consistent with classical models of volume transmission.

We then disrupted active zone organization selectively by conditionally knocking out RIM1 and RIM2 in dopamine neurons (RIM-cKO; DAT-Cre^+/-^ × RIM1^fl/fl^ × RIM2^fl/fl^). Knockout of RIM1/2 did not alter the firing rate of substantia nigra pars compacta (SNc) dopamine neurons, or striatal dopamine tissue content (Suppl. Fig S2). As previously reported, loss of RIM1/2 from dopamine neurons resulted in a near-complete loss of evoked dopamine transients measured by FSCV^20^ (Fig. 1D, F). Surprisingly, loss of spatially diffuse dopamine had little effect on D2R activation, as D2-IPSCs were largely intact and qualitatively similar to those of controls (Fig. 1D, F-G).

To confirm that D2-IPSCs in RIM-cKO mice were the result of nigrostriatal dopamine release activating D2Rs, rather than from compensation or off-target effects, we performed several pharmacological and genetic controls. D2-IPSCs were blocked by the D2R antagonist sulpiride (Suppl. Fig. S3A-B), inhibited by acute block of the vesicular monoamine transporter (VMAT2) (Suppl. Fig. S3C-D), and reduced in tandem with knocking out tyrosine hydroxylase (TH) from SNc dopamine neurons using a viral CRISPR strategy (Suppl. Fig. S3E-I). Dopamine release driving D2R activation in iSPNs also required vesicular exocytosis, as virally expressing tetanus toxin selectively in dopamine neurons of RIM-cKO mice abolished evoked D2-IPSCs (Suppl. Fig. S3J–M). The preservation of D2R activation is unlikely to reflect developmental compensation, as acute viral knockout of RIM1/2 in adult animals by midbrain delivery of Cre alongside a Cre-dependent channelrhodopsin (RIM-vKO) similarly abolished optically evoked dopamine transients while having little effect on D2-IPSCs (Fig. 1E-G; Suppl. Fig S3B). Finally, as there was no difference in D2R signaling sensitivity (EC50) or efficacy (Emax) in response to exogenous dopamine application in RIM-cKO compared to controls, upregulation of postsynaptic D2R sensitivity and coupling likely cannot account for the presence of D2R-mediated events in RIM-KO mice (Suppl. Fig. S3N–Q). Together, these results indicate that loss of RIM1/2 selectively eliminates spatially diffuse release detectable as spillover dopamine through FSCV, while leaving activation of D2Rs largely intact.

To determine whether the dissociation between spillover dopamine and point-to-point transmission is only a consequence of RIM1/2 deletion or is also a feature in wild type animals, we delivered a 1 Hz stimulation train to wild-type slices and measured frequency-dependent depression of each measure independently. Spillover dopamine transients depressed more steeply than D2-IPSCs across the train (Fig. 1G; Suppl. Fig S4), indicating that spatially diffuse dopamine does not necessarily predict receptor activation even under control conditions.

To confirm that point-to-point transmission in RIM-cKO mice engages endogenous effectors beyond exogenous GIRK channels, we next examined D2R-mediated inhibition of voltage-gated Ca²⁺ channels (VGCCs) through membrane-delimited Gβγ signaling using two-photon imaging of the Ca²⁺ indicator Fluo-5F. Electrically stimulated dopamine equally suppressed dendritic calcium influx evoked by backpropagating action potentials in iSPNs from wild-type and RIM-cKO mice, with effects in both genotypes blocked by sulpiride (Fig. 1H, Suppl. Fig S5A-C). To determine whether dopamine release in RIM-cKO mice can also engage D1 dopamine receptor (D1R)-mediated signaling, we virally expressed the genetically encoded cAMP sensor G-Flamp2^36^ selectively within direct pathway SPNs (dSPNs) and measured cAMP changes with two-photon imaging. Electrically evoked increases in G-Flamp2 activity were broadly reduced in RIM-cKO mice compared to controls, though hotspots of activation were detectable within both genotypes and blocked by application of the D1R antagonist SKF3983 (Suppl. Fig S5D-G). Thus, point-to-point dopamine transmission is capable of engaging both D2Rs and D1Rs as detected through endogenous effectors. Together, these data indicate that phasic point-to-point transmission can occur in the absence of appreciable spillover dopamine, raising the possibility that the source and regulation of dopamine underlying point-to-point transmission may be distinct from that which drives spatially diffuse release.

### Direct visualization of dopamine release hotspots in RIM-cKO mice supports point-to-point transmission

If dopamine release at specific sites is sufficient to activate dopamine receptors without generating appreciable bulk extracellular accumulation, then these signals should persist in RIM-cKO mice even as spatially diffuse release is eliminated. We first monitored subsecond dopamine dynamics *in vivo* using fiber photometry of the genetically encoded dopamine sensor GRAB-DA3m^37^ virally expressed non-selectively in the dorsolateral striatum of RIM-cKO and control mice (Fig. 2A-B). Mice periodically received a liquid reward during recordings, allowing us to distinguish spontaneous dopamine transients occurring in the absence of reward from events time-locked to reward delivery. Spontaneous transients were detectable in RIM-cKO mice (Fig. 2A-D) though reduced in frequency and amplitude relative to controls (Fig. 2A–D), consistent with prior reports^35^. Dopamine transients in both genotypes were abolished following injection of the D1R antagonist SCH23390, which blocks GRAB-DA3m activation (Fig. 2C), and absent in mice expressing control GFP only (Suppl. Fig S6). We also detected reliable reward-evoked transients in RIM cKO mice that were again reduced in amplitude compared to controls (Fig 2E). Behaviorally, we observed no difference in licking in response to reward (Suppl. Fig S6A). However, RIM cKO mice showed slower kinetics of transients associated with licking and reward delivery (Suppl. Fig S6B-C). Thus, despite the near-complete loss of spatially diffuse release detected by FSCV, subsecond dopamine transients persist *in vivo* in RIM-cKO mice, suggesting that a residual source of dopamine release evades detection by *ex vivo* bulk measurement approaches.

**Figure 2:**
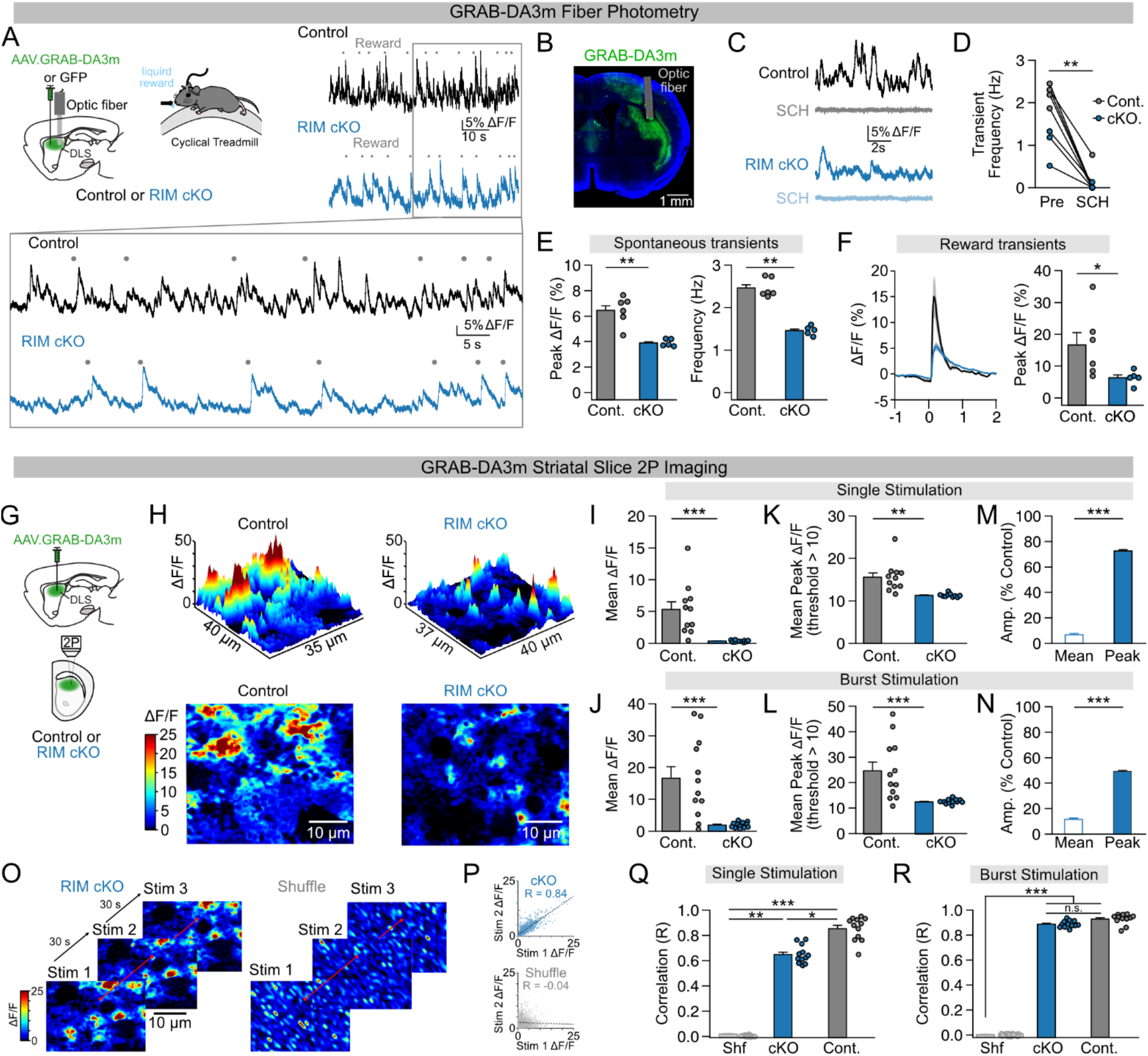
Imaging dopamine in RIM cKO mice reveals localized hotspots of release consistent with point-to-point transmission. A. Schematic of the approach for measuring bulk GRAB-DA3m activity in mice periodically receiving a liquid reward (top left). Example photometry traces from control and RIM cKO mice (top right), with expanded window below. Grey dots represent reward delivery (solenoid opening). B. Example image of GRAB-DA3m expression with the optical fiber track highlighted in grey. C. Example photometry traces from control and RIM cKO mice before and after administration of the D1R antagonist SCH23390. D. Quantification of the effect of SCH23390 on transient frequency (** p<0.01). Wilcoxon signed-rank test. E. Quantification of the mean amplitude (left, ** p<0.01) and frequency (right, ** p<0.01) of spontaneous transients in control and RIM cKO mice. Wilcoxon rank-sum tests. F. Quantification of the mean amplitude of reward evoked transients in control and RIM cKO mice (* p<0.05) Wilcoxon rank-sum test. G. Schematic of method used to map dopamine release in striatal slices using 2-photon imaging of GRAB-DA3m. H. Example images of electrically evoked GRAB-DA3m activity in slices from control and RIM cKO mice. I. Quantification of the mean GRAB-DA3m response for a single stimulation in slices from control (Cont.) and RIM cKO (cKO) mice (*** p<0.001). Wilcoxon rank-sum test. J. Quantification of the mean GRAB-DA3m response for a burst stimulation (40 Hz, 5 pulses) in slices from control and RIM cKO mice (*** p<0.001). Wilcoxon rank-sum test. K. Quantification of the mean response within hotspots (Mean Peak) for a single stimulation. Hotspots were defined as regions with an amplitude greater than 10 ΔF/F (** p<0.01). Wilcoxon rank-sum test. L. Quantification as in (K), but for burst stimulation (** p<0.01). Wilcoxon rank-sum test. M. Relative change in mean GRAB-DA3m response (Mean) and mean hotspot amplitude (Peak) for RIM cKO mice compared to controls (*** p<0.001). Data is for single pulse stimulation. Wilcoxon rank-sum test. N. Quantification as in (M), but for burst stimulation. Wilcoxon rank-sum test. O. Depiction of approach for correlating GRAB-DA3m responses across stimulation bouts with original and shuffled data. P. Example correlation plots for original and shuffled data across the first and second stimulus trial. Q. Quantification of the mean correlation across stimulation trials for a single pulse. Kruskal-Wallis test (*** p<0.001) followed by Tukey’s post hoc test (Shf vs cKO: ** p<0.01, Shf vs Cont.: *** p<0.001, cKO vs Cont.: * p<0.05). R. Quantification of the mean correlation across stimulation trials for a burst stimulus. Kruskal-Wallis test followed by Tukey’s post hoc test. Kruskal-Wallis test (*** p<0.001) followed by Tukey’s post hoc test (Shf vs cKO: *** p<0.001, Shf vs Cont.: *** p<0.001, cKO vs Cont.: ns p>0.05). Summary data are the mean ± SEM.

Having detected phasic dopamine activity *in vivo*, we next used two-photon imaging of GRAB-DA3m non-selectively expressed in *ex vivo* dorsolateral striatal slices to examine if spatially discrete dopamine release hotspots are present in RIM-cKO mice (Fig. 2G). Electrical stimulation increased sensor fluorescence in slices from both control and RIM-cKO animals (Fig 2H). Consistent with FSCV measurements, the spatially averaged sensor response across the field of view was dramatically reduced in RIM-cKO compared to controls following both single and burst stimulation (Fig. 2H-J), confirming the loss of spatially diffuse release. Given that point-to-point D2R activation in ex vivo slices requires high local dopamine concentrations, we next examined if hotspots of high sensor activation could be detected in RIM cKO mice (Suppl. Fig S7). Evoked increases in GRAB-DA3m fluorescence (ΔF/F) were thresholded at different detection levels to identify putative hotspots (Suppl. Fig S7). We first chose a threshold of 10 ΔF/F, as this corresponds to the averaged response of the sensor to dopamine concentrations of ∼1 μM^37^. At these concentrations, D2R-mediated Gβγ signaling would be expected to be engaged under physiological conditions, suggesting these sites may preferentially reflect point-to-point release events^28^. Regions exceeding this threshold were consistently detectable in slices from both wild-type and RIM-cKO mice (Fig. 2H, K-L) and were absent in the presence of SCH23390 (Suppl. Fig. S8). Mean sensor activation within hotspots was partially reduced in RIM-cKO (Fig. 2K-L). Paralleling the partial reduction in point-to-point transmission despite the loss of spatially diffuse release (Fig. 1D), we observed only a partial reduction in the hotspot amplitude which was in contrast to the near complete reduction in the averaged whole frame response (Fig. 2M-N). Similar results were obtained across thresholds spanning the dynamic range of GRAB-DA3m (Suppl. Fig S9) and in experiments using the alternate sensor dLight1.3b^26^ expressed in striatal dopamine axons (Suppl Fig. S10).

If sparse, point-to-point dopamine transmission occurs, we would expect its location to be consistent across repeated stimulation trials (Fig. 2O). Indeed, GRAB-DA3m fluorescence transients were highly spatially correlated across trials in slices from both control and RIM cKO mice, compared to shuffled data (Fig. 2O-R). Interestingly, the degree of correlation in the RIM cKO group was lower using single stimulation compared to burst stimuli (40 Hz, 5 pulses), potentially reflecting lower dopamine release probability following the loss of RIM1/2 (Fig. 2Q-R). Together, these findings indicate that spatially localized dopamine release is present in RIM-cKO mice, albeit at lower levels than in slices from control animals, consistent with local transmission remaining largely intact in the absence of spatially diffuse release.

### Loss of spillover dopamine alters SPN excitability but not spine density

Given the persistence of dopamine release hotspots and phasic D2R activation despite the near-complete loss of spatially diffuse release in RIM-cKO mice, we asked whether these two modes of dopamine signaling differentially shape SPN physiology. As dopaminergic lesions are known to induce both homeostatic adaptations in the intrinsic excitability as well as reductions in the spine density of dSPNs and iSPNs^38–41^, we leveraged the RIM-cKO model to determine which of these adaptations are driven by the loss of spatially diffuse release and which may be sustained by residual point-to-point transmission.

To assess intrinsic excitability, we measured current-induced action potential firing in whole-cell recordings of dSPNs and iSPNs in slices from wild-type and RIM-cKO mice (Fig. 3A-H). In RIM-cKO mice, dSPNs exhibited a leftward shift in excitability, reflected in reduced rheobase and increased input resistance (Fig. 3B-D). Conversely, iSPNs showed a rightward shift in excitability reflected in increased rheobase but no change in input resistance (Fig. 3F–H; Suppl. Fig. S11A-C). This bidirectional pattern is consistent with the homeostatic adaptations in SPN excitability observed following dopaminergic lesions^40^.

**Figure 3:**
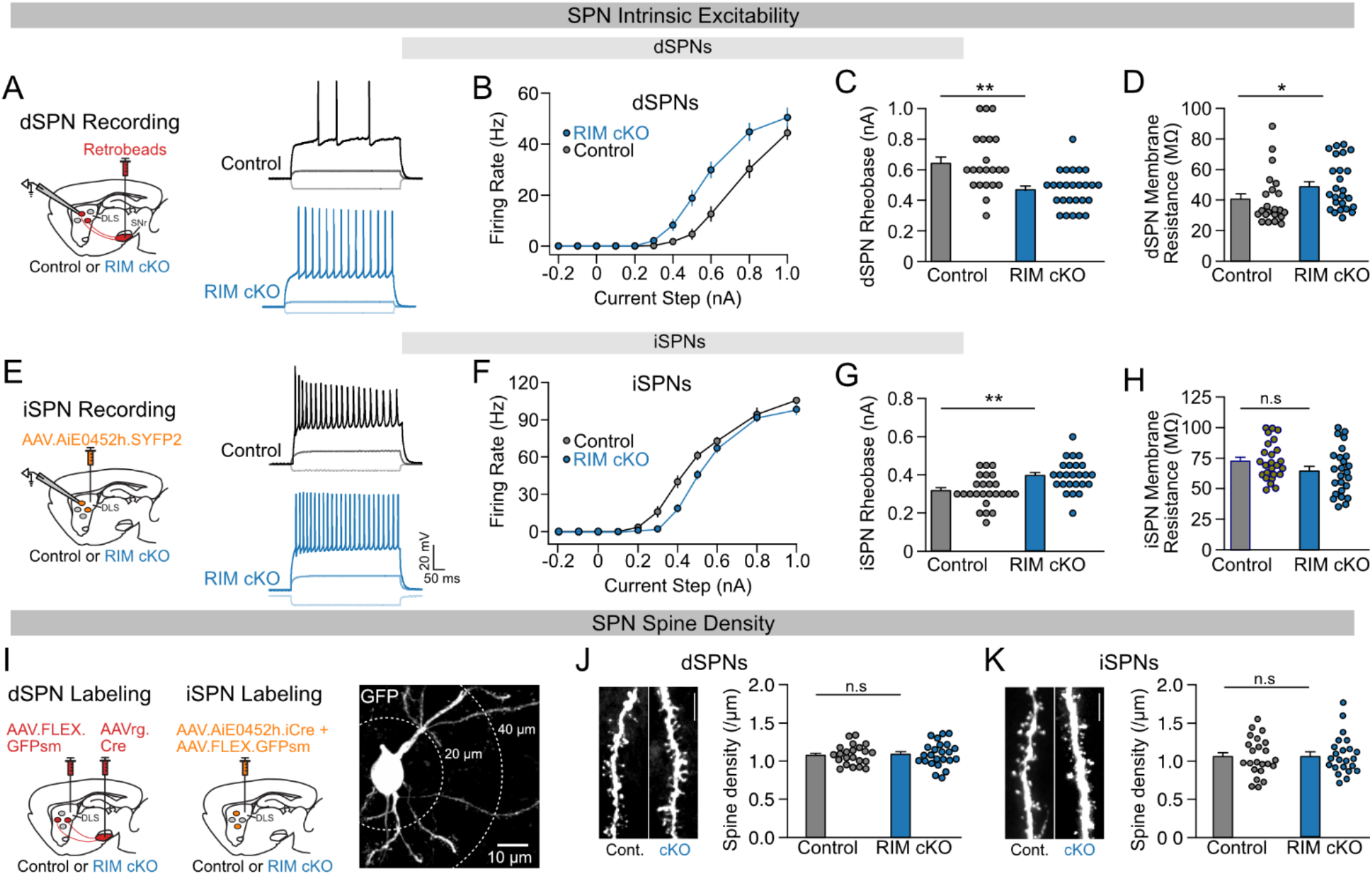
Loss of spatially diffuse dopamine release alters striatal circuit excitability, but not SPN spine density. A. Schematic showing the use of retrobeads injected in the SNr to label dSPNs in control and RIM cKO mice (left). Example traces of current-induced spiking in dSPNs from control and RIM cKO mice. B. Action potential firing rate responses for dSPNs from control and RIM cKO mice. C. Quantification of rheobase in dSPNs from control and RIM cKO mice (** p<0.01). Wilcoxon rank-sum test. D. Quantification of the average membrane resistance of dSPNs from control and RIM cKO mice (* p<0.05). Wilcoxon rank-sum test. E. Schematic showing the use of an enhancer specific AAV to label iSPNs in control and RIM cKO mice (left). Example traces of current-induced spiking in iSPNs from control and RIM cKO mice. F. Action potential firing rate responses for iSPNs from control and RIM cKO mice. G. Quantification of rheobase in iSPNs from control and RIM cKO mice (** p<0.01). Wilcoxon rank-sum test. H. Quantification of the average membrane resistance of dSPNs from control and RIM cKO mice (ns p>0.05). Wilcoxon rank-sum test. I. Schematic depicting the approach for quantifying spine density in SPNs from control and RIM cKO mice (left). iSPNs and dSPNs were selectively labeled with fluorescent reporter using similar approaches to those described in (A) and (E). Example image of a GFP^+^ labeled SPN. J. Example images of dendrites from dSPNs from control (Cont.) and RIM cKO (cKO) mice (left, scale bar = 5 µm). Quantification of dendritic spine density in dSPNs from control and RIM cKO mice (right, ns p>0.05). Wilcoxon rank-sum test. K. Example images of dendrites from iSPNs from control (Cont.) and RIM cKO (cKO) mice (left, scale bar = 5 µm). Quantification of dendritic spine density in iSPNs from control and RIM cKO mice (right, ns p>0.05). Wilcoxon rank-sum test. Summary data are mean ± SEM.

We next assessed dendritic spine density in both proximal and distal dendrites with cell-type specific confocal imaging of fluorescently labeled dSPNs and iSPNs (Fig. 3I-K). In contrast to dopaminergic lesion models, in which spine density is reduced in both SPN populations, RIM-cKO mice showed no difference in spine density in either dSPNs or iSPNs (Fig. 3J–K, Suppl Fig.11D-E). Together, these results indicate that spatially diffuse release is required for maintaining SPN intrinsic excitability but appears dispensable for dendritic spine density, suggesting that point-to-point transmission and local activation of receptors may be sufficient to sustain the structural integrity of SPN spines.

### Spatially diffuse dopamine promotes locomotion but is not required for motor learning

Intact dopamine transmission within the dorsolateral striatum is essential for motor output and motor learning, yet how dopamine signaling within this region independently regulates both aspects of motor behavior remains poorly understood. Having found that loss of spatially diffuse release recapitulates some, but not all, of the SPN adaptations observed following dopaminergic lesions, we next asked whether spatially diffuse release and point-to-point transmission differentially contribute to normal motor behavior as well.

We first used spontaneous locomotion of animals in the open field as an assay of gross motor output (Fig. 4A). In addition to comparing RIM-cKO and RIM-vKO mice to control animals, we utilized mice treated with a low dose of reserpine (0.5 mg/kg). These animals served as a positive control as they show similar partial reductions in both spatially diffuse and point-to-point dopamine transmission (Figure 1C). As expected, reserpine-treated mice exhibited reductions in mean velocity, time mobile, and average peak velocity compared to controls (Fig. 4B–D), similar to mice unilaterally treated with 6-OHDA (Suppl. Fig S12A-B). Similar reductions in locomotion across all three measures were observed in both RIM-cKO and RIM-vKO mice (Fig. 4B–D), indicating that loss of spatially diffuse dopamine is sufficient to impair motor output.

**Figure 4:**
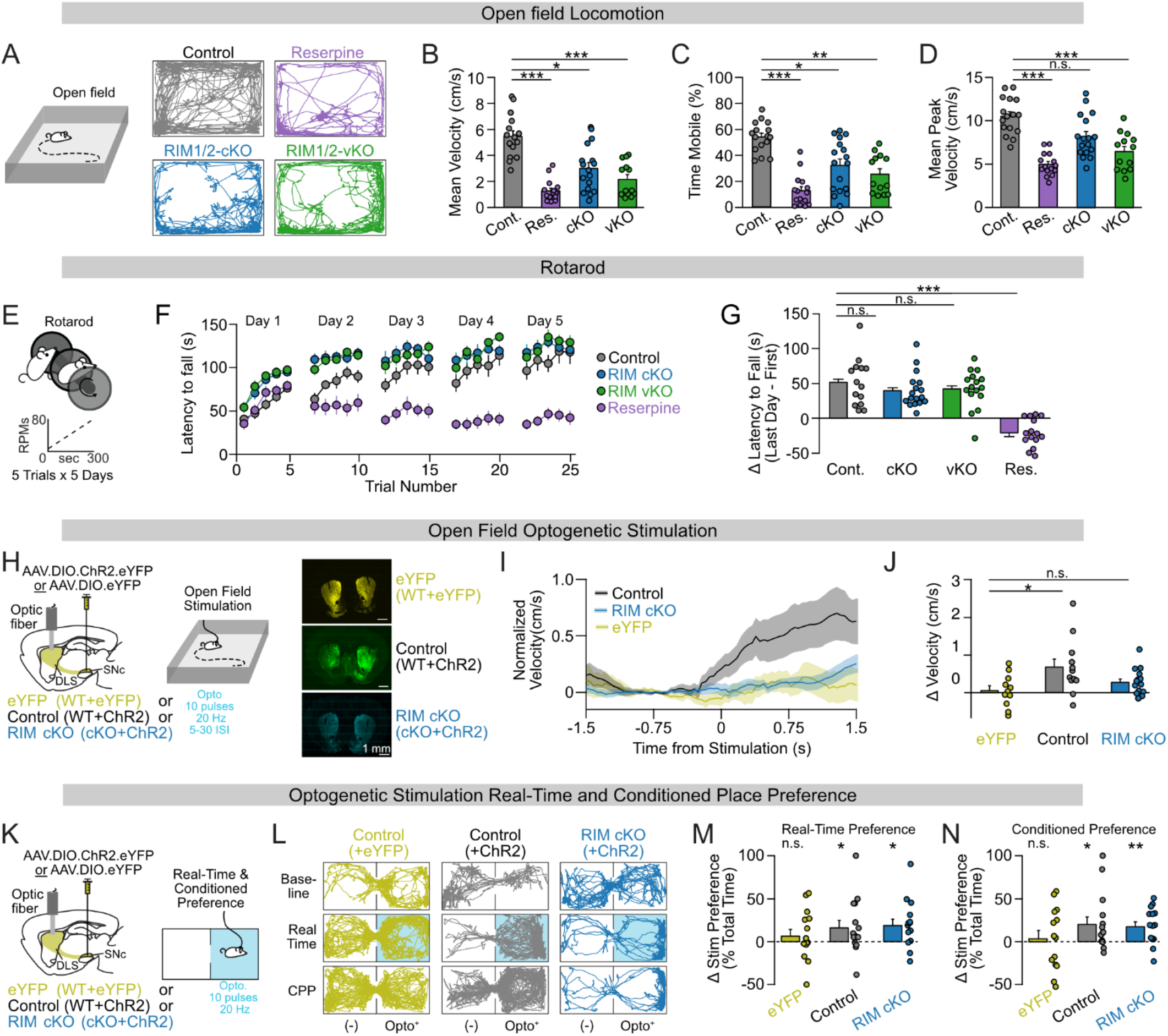
Spatially diffuse dopamine promotes motor output while point-to-point transmission supports learning. A. Schematic of the open field assay with example position plots from control, reserpine-treated, RIM cKO, and RIM vKO mice. B. Quantification of mean velocity for control (Cont.), reserpine-treated (Res.), RIM cKO (cKO) and RIM vKO (vKO) mice. Kruskal-Wallis test (*** p<0.001) followed by Dunnet’s post hoc test (Cont vs. Res.: *** p<0.001, Cont. vs. cKO: * p<0.05, Cont. vs vKO: *** p<0.001). C. Quantification of percent time mobile for the groups listed in (B). Kruskal-Wallis test (*** p<0.001) followed by Dunnet’s post hoc test (Cont vs. Res.: *** p<0.001, Cont. vs. cKO: * p<0.05, Cont. vs vKO: ** p<0.01). D. Quantification of percent mean peak velocity for the groups listed in (B). Kruskal-Wallis test (*** p<0.001) followed by Dunnet’s post hoc test (Cont vs. Res.: *** p<0.001, Cont. vs. cKO: ns p>0.05, Cont. vs vKO: *** p<0.001). E. Schematic of the accelerating rotarod assay. F. Rotarod performance (latency to fall) across trials and days for the groups listed in (B). G. Quantification of the change in performance for data shown in (F). Kruskal-Wallis test (*** p<0.001) followed by Dunnet’s post hoc test (Cont vs. cKO.: ns p>0.05, Cont. vs. vKO: ns p>0.05, Cont. vs Res.: *** p<0.001). H. Schematic of approach for optogenetically stimulating nigrostriatal dopamine axons in the open field (left). Example images of viral expression and fiber tracks eYFP, control, and RIM cKO mice (right). I. Changes in velocity following light delivery during periods of immobility (H). J. Quantification of the data shown in (I). Kruskal-Wallis (* p<0.05) followed by Tukey’s post hoc test (eYFP vs. Control: * p<0.05, eYFP vs RIM cKO: ns p>0.05, Control vs. RIM cKO: ns p>0.05). K. Schematic of optogenetic real-time and conditioned place preference (CPP) assays. L. Example position plots from eYFP, Control, and RIM cKO mice during the baseline, real-time, and CPP sessions. M. Quantification of the change in preference for the light-paired chamber during real-time place preference assay for eYFP (ns p>0.05), control (* p<0.05), and RIM cKO mice (* p<0.05). Wilcoxon signed-rank tests. N. Quantification in the change in preference for the light-paired chamber during the CPP assay for eYFP (ns p>0.05), control (* p<0.05), and RIM cKO mice (** p<0.01). Wilcoxon signed-rank tests. Summary data are mean ± SEM.

We next asked how loss of spatially diffuse dopamine impacts motor learning using the accelerating rotarod assay, a task in which mice learn to remain on an accelerating wheel for increasing durations across days (Fig. 4E). While control mice improved across days of training (Fig. 4E-G), both reserpine and 6-OHDA treated mice failed to show improvements across days (Fig. 4E-G, Suppl. Fig. S12C-E). In contrast, RIM-cKO mice improved both within and across sessions at rates indistinguishable from wild-type animals (Fig. 4F-G). Preserved motor learning in the absence of spatially diffuse release was not a consequence of developmental compensation, as similar results were obtained in RIM-vKO mice (Fig. 4F-G). Notably, all groups showed similar learning within days (Suppl. Fig. S12F-G), consistent with dopamine signaling mediated consolidation of learning across days ^42^. These data suggest that point-to-point transmission is sufficient to support motor learning in the near absence of spatially diffuse dopamine.

To more causally test whether spatially diffuse release is required for dopamine’s modulatory effects on movement, we virally expressed ChR2 selectively in dopamine neurons of wild-type and RIM-cKO mice and implanted bilateral optical fibers in the dorsolateral striatum (Fig. 4H) to examine the effect of optical stimulation in the open field, which was previously shown to promote movement initiation^3,4^. Mice expressing eYFP in dopamine neurons served as controls for viral expression, fiber implantation, and light delivery. Randomized delivery of short bursts of optogenetic stimulation favored movement initiation in ChR2-expressing control mice as indicated by an increase in velocity, but failed to do so in RIM-cKO mice or eYFP controls (Fig. 4I–J; Suppl. Fig S12H-J), indicating that spatially diffuse release is required for the modulatory effects of DA on movement initiation.

Phasic dopamine release is essential for associative learning and for the formation of contextual reward memories. Whether point-to-point transmission is sufficient to support these functions in the absence of spatially diffuse release remains unknown. We next tested ChR2- and eYFP-expressing animals in a real-time and conditioned place preference paradigm in which one compartment of a two-chambered box was paired with optical stimulation of dopamine terminals (Fig. 4K). ChR2-expressing control and RIM-cKO mice both increased their preference for the light-paired compartment during real-time place preference assays. This preference was maintained during a subsequent test session without light stimulation, indicative of a learned association (Fig. 4L-N; Suppl. Fig. S12K-L). Mice expressing eYFP showed no change in preference in either session (Fig. 4L-N; Suppl. Fig. S12K-L). No difference in crossings was observed across groups (Suppl. Fig 12M). Together, these results indicate that spatially diffuse release and point-to-point transmission make distinct contributions to behavior, with spatially diffuse release mediating the effects of dopamine on locomotion and point-to-point transmission supporting associative and motor learning.

## Discussion

Here we sought to understand how distinct spatiotemporal modes of dopamine transmission shape striatal function and motor behavior. We found that spatially diffuse release, which underlies the volume transmission framework, appears necessary for maintaining SPN excitability and for normal locomotion. Point-to-point transmission, assessed through direct measurement of postsynaptic dopamine receptor activation, persisted in the near-complete absence of spatially diffuse release and appeared sufficient to preserve dendritic spine density and support both associative and motor learning. Together, these findings suggest that the geometry of dopamine release may be a key determinant of how dopaminergic signals are decoded by striatal circuits to regulate distinct aspects of behavior.

The relationship between the temporal structure of dopamine release and its influence on motor output has been a subject of debate. Prior work has shown that dopamine neuron activity increases prior to movement onset, and that phasic activation of dopamine axons is sufficient to promote movement initiation^3–6^. However, more recent findings suggest that tonic dopamine concentrations are sufficient for normal locomotion, with phasic release being largely dispensable^35,43,44^. Our results suggest that both the temporal and spatial structure of dopamine transmission are critical determinants of its impact on movement initiation, with locomotion requiring release patterns that generate spatially diffuse dopamine in the extracellular space. This is consistent with the observation that levodopa restores motor function by broadly elevating dopaminergic tone without reinstating phasic transients^43^. The kinetics of spatially diffuse dopamine release, however, suggest that its influence on movement operates through modulation of movement probability and locomotor vigor rather than direct moment-to-moment gating of motor output. This view is consistent with our finding that point-to-point transmission, while operating on faster timescales, is insufficient to directly promote locomotion. Prior observations that supraphysiological dopamine transients are required for optogenetic stimulation to enhance motor output, but not to reinforce behavior^5^, may similarly reflect the spatial distribution of dopamine release, with higher-intensity stimulation generating the spatially diffuse dopamine required to enhance ongoing locomotion.

Our findings also provide insight into how dopamine contributes to motor learning. The preservation of associative and motor learning in RIM-cKO mice indicates that point-to-point transmission is sufficient to gate the synaptic plasticity underlying skill acquisition. Dopamine is a critical determinant of corticostriatal synaptic plasticity^45–47^ involved in refining precisely timed movements during skill acquisition^48,49^. The narrow time window of dopamine-dependent structural plasticity^50,51^ and the clustered potentiation of inputs along SPN dendrites^49^ that occur during motor learning suggest that this gating is both temporally and spatially precise. By signaling locally and tracking dopamine neuron firing more faithfully than high-probability release sites, RIM-independent point-to-point transmission may provide a mechanism by which dopamine contributes to the precise modulation of corticostriatal synaptic strength during learning.

The differential dependence of spatially diffuse release and point-to-point transmission on RIM1/2 suggests a mechanism by which the molecular composition of individual release sites^20,52–54^ may determine the spatiotemporal profile of dopamine signaling. As RIM proteins are required for high-probability vesicular release^55–57^, their presence at a subset of release sites would be expected to produce sufficient synchronous release to overwhelm reuptake and generate bulk extracellular dopamine accumulation. Such release would show strong frequency-dependent depression, reducing its ability to follow individual action potentials. Conversely, release sites with reduced or absent RIM dependence would exhibit lower release probability, making them more resistant to depression and enabling more faithful tracking of dopamine neuron firing across tonic and phasic activity patterns. Such sites could generate locally high dopamine concentrations capable of activating receptors without contributing to bulk extracellular accumulation.

An important question raised by these findings is how presynaptic release site heterogeneity relates to the postsynaptic organization of dopamine receptors. At classical glutamatergic and GABAergic synapses, release properties can differ across terminals from the same neuron depending on the identity of the postsynaptic target, providing a mechanism for target-specific transmission^58–60^. Whether analogous heterogeneity exists at neuromodulatory synapses has not been established, though recent anatomical evidence suggests that dopamine axons show a preferential association with receptor-expressing dendrites, with release-capable varicosities clustering near postsynaptic targets in a synapse-like organization^61^. Our findings provide functional evidence consistent with this view, suggesting that postsynaptic target identity may determine the mode of dopamine transmission received. It is important to note that while our data establish a functional dissociation between spatially diffuse extracellular dopamine and local point-to-point receptor activation, they do not by themselves resolve the molecular basis of this dissociation. The observation that partial depletion of vesicular dopamine content with reserpine reduces both FSCV signals and D2R-mediated transmission in parallel argues against a simple difference in sensitivity between the two readouts, supporting the view that the RIM cKO phenotype reflects a genuine functional dissociation rather than a detection threshold effect. However, the molecular basis of this functional dissociation and whether it reflects two distinct release modalities remain to be determined.

The distinct behavioral effects of different modes of dopamine transmission suggest that downstream SPNs deconvolve these signals. In line with this prediction, we find that changes in SPN excitability and spine density differentially depend on spatially diffuse and point-to-point transmission. The bidirectional excitability changes in dSPNs and iSPNs of RIM-cKO mice provide a cellular mechanism for the locomotor deficit: reduced spatially diffuse dopamine decreases dopaminergic gain on the direct/indirect pathway balance, reducing net striatal output and locomotor drive. Likewise, the preservation of SPN spine density in RIM-cKO mice suggests that point-to-point transmission supports motor learning by maintaining plasticity mechanisms at excitatory synapses. These processes may be spatially segregated within cellular compartments or potentially depend on distinct secondary messenger cascades that also differ in the spatiotemporal dynamics^28^.

More broadly, our findings carry important implications for disease and for neuromodulatory transmission in general. Impaired dopamine signaling is implicated in disorders such as Parkinson’s disease, schizophrenia, ADHD, Tourette’s syndrome, and addiction. Despite the stark difference in symptomology across these diseases, dopaminergic dysfunction is often portrayed along a single axis as either a paucity or excess. By incorporating the geometry of dopamine release, our results expand the axes on which dopaminergic dysfunction can be understood and linked to alterations in behavior. They also raise the question of whether other neuromodulatory systems such as the serotonergic and noradrenergic systems similarly multiplex information through different forms of release. The answer likely carries implications for numerous other neurological disorders.

## Methods

### Experimental Animals

All animal procedures were conducted in accordance with protocols approved by the Institutional Animal Care and Use Committee (IACUC) at the University of Colorado Anschutz School of Medicine, McGill University and New York University. Animals were group or singly housed in a temperature and humidity-controlled environment on a 12-hour light-dark cycle and allowed ad libitum access to food and water. Male and female mice lacking RIM1 and RIM2 proteins in dopamine neurons (RIM-cKO mice) were generated by crossing RIM1 (RRID:IMSR_JAX: 015832, Rims1tm3Sud/J) and RIM2 (RRID:IMSR_JAX: 015833, Rims2tm1.1Sud/J) double floxed mice with heterozygous DAT-IRES-Cre mice (Kaeser et al., 2011) and were initially provided by Pascal Kaeser. Control animals included DAT-Cre negative littermates or DAT-Cre-positive mice (DAT-Cre^+/-^ x RIM1/2^wt/wt^) from the same background when selective genetic access to dopamine neurons was required. All experiments were performed in adult animals (> P42).

### Stereotaxic Injections and Optical Cannula Implants

Adult mice were anesthetized with 2% isoflurane (induction and maintenance) and mounted on a stereotaxic frame (Kopf Instruments). Injections were performed using a pulled glass pipette filled with mineral oil and a Nanoject III (Drummond Scientific) set to an injection rate set to 2nL/sec. Following the injection, the pipette was left in place for 5 minutes, then slowly removed. Mice were given carprofen (i.p., 5 mg/kg) twenty minutes before surgery completion and allowed to recover in a cage on a heating pad until awake after the scalp was sealed with Vetbond (1469SB, 3M). Carprofen injections were repeated the following day, after which mice were monitored daily until they fully recovered. For unilateral lesions with 6-OHDA, mice were kept in a cage on a heating pad and given daily injections of warmed sterile saline (1 mL, subcutaneous) for one to two weeks. During this period, they were provided with high calorie diet gel (72-06-5022, Clear H20) and their weight monitored daily.

All stereotaxic coordinates were determined using the distance from bregma (anteroposterior and mediolateral) and the dural surface (dorsoventral) in millimeters. The following coordinates were used: Dorsolateral striatum (DLS, AP: 0.8, ML: 2.2, DV: −2.8), Globus Pallidus externa (GPe, AP: −0.29, ML: 1.75, DV: −3.58), Substantia Nigra pars compacta (SNc, AP: −3.1, ML: 1.25, DV: −4.7), and Medial Forebrain Bundle (MFB, AP: −1.2, ML: 1.3, DV: −4.75). See Table 1.1 for a list of viruses and reagents with injection site, volume, dilution, and identifying number.

**Table 1.1.**

| **Experiment** | **Virus** | **Site** | **Volume** | **Dilution** | **Provider** | **ID** |
| --- | --- | --- | --- | --- | --- | --- |
| <u>GIRK Recordings</u> | AAV9-Syn-tdTomato-T2A-mGIRK2 | DLS | 400 nL | 1:3 | UPenn Viral Core | V3992 |
| <u>Viral RIM Knockout</u> | AAV5-Ef1a-Cre-mcherry and |  |  | 1:3 | UNC Viral Core | 55632 |
|  | AAV5-EF1a-DIO-hChR2(H134R)-eYFP | SNc | 400 nL | 1:3 | Addgene | 20298-AAV5 |
| <u>TH Knockout</u> | AAV1-CMV-FLEX-SaCas9-U6-sgTh | SNc | 400 nL | 1:1 | Addgene | 159901 |
| <u>Tetanus Toxin Expression</u> | AAV5-hSyn-DIO-Halotag-2A-TeNT or | SNc | 400 nL | NA | Kennedy Lab | NA |
|  | AAV9-pCAG-FLEX-EGFP | SNc | 400 nL | 1:9 | Addgene | 51502-AAV9 |
| <u>6-OHDA Lesions</u> | 6-hydroxydopamine (6-OHDA) | MFB | 1000 nL | 4µg/µL | Sigma-Aldrich | 162957 |
| <u>Striatal DA Imaging</u> | AAV1-hSyn-GRAB-DA3m or | DLS | 400 nL | 1:3 | Addgene | 208698-AAV1 |
|  | AAV9-Syn-dLight1.3b | DLS | 400 nL | 1:7 | Addgene | 135762-AAV9 |
| <u>Axonal DA Imaging</u> | AAV5-FLEX-CAG-dLight1.3b and |  |  | 1:7 | Vigene | 6833 |
|  | pAAV5-CAG-tdTomato | SNc | 400 nL | 1:79 | Addgene | 59462-AAV5 |
| <u>Intrinsic Excitability</u> | AAV9-AiE0452h-minBG-SYFP2 | DLS | 400 nL | 1:6 | Addgene | 224173 |
| <u>Retrograde labeling</u> | AAVrg-hSyn-Cre-P2A-dTomato or | Gpe | 200 µL | NA | Addgene | 107738-AAVrg |
|  | AAVrg-hSyn-Cre-P2A-dTomato and | SNr | 400 nL | NA | Addgene | 107738-AAVrg |
|  | AAV1-CAG-FLEX-GFPsm_myc | DLS | 400 nL | 1:19 | Addgene | 98927-AAV1 |
|  | Red Retrobeads or | Gpe | 200 nL | 1:3 | Lumafluor | NA |
|  | Red Retrobeads | SNr | 400 nL | 1:3 | Lumafluor | NA |
| <u>Optogenetics</u> | AAV5-EF1a-DIO-hChR2(H134R)-eYFP | SNc | 400 nL | 1:3 | Addgene | 20298-AAV5 |
|  | or AAV5-EF1a-DIO-EYFP | SNc | 400 nL | 1:1 | Addgene | 27056-AAV5 |
| <u>Fiber Photometry</u> | AAV9-hSyn-GRAB-gDA3m or | DLS | 400 nL | 1:10 | Biohippo | PT-4720 |
|  | AAV1-CMV-PI.EGFP-WPRE-bGH | DLS | 400 nL | 1:10 | Addgene | 105530-AAV1 |
| <u>cAMP Imaging</u> | pAAV-AiE0779m_3xC2-minBG-iCre(R297T) and |  |  | 1:7 | Addgene | 199776-PHPeB |
|  | AAV2/9-hSyn-DIO-opt-GFlamp2 | DLS | 400 nL | 1:7 | BCH Viral Core |  |
| <u>VGCCs Activity</u> | AAV9-AiE0452h-minBG-SYFP2 | DLS | 400 nL | 1:3 | Addgene | 224173 |

For *in vivo* optogenetics, homemade optical canulae of custom length were made using optical fiber (FT200EMT, Thorlabs), ceramic ferrules (CFLC230-10, Thorlabs), and epoxy (G14250, Thorlabs). Fibers were cut flat with a length of 4-5mm and polished to a minimum of 80% efficiency. Commercial optical canulae (MFC_200/250-0.66_4.5mm_MF1.25_FLT, Doric) were used for fiber photometry experiments. Following viral injections, fibers were bilaterally implanted within the dorsal striatum (AP:0.8 ML: 2.2 DV: −2.8) and affixed to the skull using C&B Metabond (S380, Parkell) and Ortho-Jet dental cement (1334CLR, Lang). Vetbond was used to ensure the scalp was secured around the base of the cement.

For fiber photometry experiments, (AAV) encoding either the green GRAB-DA3m or GFP was injected in the DLS (AP: 0.4, ML: 2.4 mm, DV: −2.5 mm) using a pulled glass injection micropipette (Wiretrol II; Drummond) connected to a syringe pump (Legato 111; KD scientific) fitted with a Hamilton syringe (Gastight 1701N) via PE tubing filled with mineral oil. A 4-mm-long optical cannula (400-μm core, 0.48 numerical aperture; RWD Life Science) was subsequently implanted 0.2 mm above the injection site. For head-fixing, a custom titanium headpost was attached to the skull in the horizontal plane posterior to lambda using C&B Metabond (Parkel).

### Slice Preparation

Mice were deeply anesthetized with isoflurane and transcardially perfused with ice-cold cutting solution containing (in mM): 75 NaCl, 2.5 KCl, 6 MgCl2, 0.1 CaCl2, 1.2 NaH2PO4, 25 NaHCO3, 2.5 D-glucose, 50 sucrose (bubbled with 5% CO2 in O2). The brain was quickly removed, blocked, and affixed to a cutting block with glue. Coronal striatal or midbrain sections (240 µm thick) were cut in the same ice-cold cutting solution using a vibratome (VT1200S, Leica). After cutting, slices were placed in 32°C artificial cerebrospinal fluid (ACSF) containing (in mM): 126 NaCl, 2.5 KCl, 1.2 MgCl2, 2.0 CaCl2, 1.2 NaH2PO4, 21.4 NaHCO3, 11.1 D-glucose and 0.01 MK-801. ACSF was continuously bubbled with carbogen (5% carbon dioxide, 95% oxygen) and slices were left to recover for a minimum of 30 minutes.

### Slice Electrophysiology

For whole-cell patch clamp recordings, slices were transferred to a recording chamber where they were continuously superfused at a rate of ∼2mL/min with 32°C ACSF bubbled with carbogen. Cells imaged using an Olympus BX51WI microscope with a 40x objective. Glass pipettes (o.d. 1.5 mm, 2-4 MΩ, Word Precision Instruments) pulled on a vertical puller (P-10, Narishige) were used to patch SPNs in the dorsolateral striatum. An Axopatch 200B amplifier (Molecular Devices) and Axograph (Axograph Scientific) were used to acquire data that was sampled at 10 kHz and filtered at 5 kHz.

To isolate electrically or optically evoked D2R-mediated IPSCs, ACSF contained the following drugs (in µM): 10 DNQX, 100 picrotoxin, 0.3 CGP 35348, 1.0 SCH23390, 1.0 DHβE, and 0.3 scopolamine. Internal solution contained the following (in mM): 135 potassium gluconate, 10 HEPES, 0.1 CaCl2, 2 MgCl2, 10 BAPTA, as well as (in mg/mL) 1 ATP, 0.1 GTP, and 1.5 phosphocreatine. SPNs were held in voltage clamp at −60 mV and D2R-IPSCs elicited with electrical stimulation (1 pulse, 40µA, 0.7 ms) delivered from a stimulus isolator (A365, Word Precision Instruments) using a monopolar glass stimulating electrode inserted into the slice approximately one field of view away. A wide-field LED (Thorlabs) was used to deliver brief pulses of blue light (470 nm, 2 ms, 4 mW) for optically evoked D2R-IPSCs. In all experiments, D2R-IPSCs were evoked regularly every 1-2 minutes and a brief −2mV voltage step prior to stimulation was used to monitor series resistance. Custom Matlab code (2019a, Mathworks) was used to quantify the amplitude of D2R-IPSCs, which was calculated by finding the peak current response within 1 second of stimulation and taking the difference from the baseline current. A separate period from the baseline was used as a ‘mock’ response to estimate the contribution of noise fluctuations and this value was subtracted from the peak amplitude.

For measuring D2R-mediated currents in response to exogenous dopamine application, 10 µM cocaine was included in the ACSF for the duration of the recording to block dopamine reuptake. Dopamine stock solutions were dissolved in 200 µM ascorbic acid to prevent oxidation. Following establishment of a steady baseline holding current, dopamine was bath applied until steady state current was achieved, then washed out. Series resistance was monitored using brief voltage steps during the course of the recording. Dopamine-induced current changes were manually quantified using Igor Pro (Version 6, WaveMetrics).

In general, iSPNs were identified by the presence of a D2R-IPSC in response to a single pulse (or 25 Hz burst stimulus if no response was detected). We monitored response rates in RIM-cKO and RIM-vKO mice and found no difference from their respective controls. For animals treated with 6-OHDA or viral expression of tetanus toxin, we did not distinguish between iSPNs and dSPNs as no D2R-IPSCs were ever detected. For both experiments, we included positive controls (exogenous dopamine wash-on or recordings from contralateral hemispheres) to ensure that the lack of detectable D2R-IPSCs did not result from technical issues.

Measurement of SPN intrinsic excitability was performed under similar conditions with the following modifications. External ACSF solution contained the following drugs (in µM): 10 DNQX, 100 picrotoxin, 0.3 CGP 35348, 1.0 SCH23390, 1.0 DHβE, 0.3 scopolamine, and 0.5 sulpiride. Internal solution contained the following (in mM): 115 potassium methyl sulphate, 20 sodium chloride, 1.5 magnesium chloride, 10 HEPES potassium, 10 BAPTA as well as (in mg/mL) 1 ATP, 0.1 GTP, and 1.5 phosphocreatine. Cells were held in current clamp at their resting potential and allowed to dialyze. Spiking was induced by injecting 500 ms current steps ranging in amplitude from −200 to 1200 pA. Custom Matlab code was used to identify spikes based on a voltage threshold and the firing calculated as the number of spikes divided by the duration of the current injection.

Cell-attached recordings of SNc dopamine neurons were performed under the same conditions as for measuring SPN excitability, but with external ACSF used as internal solution. Custom Matlab code was again used to identify spikes and firing rates were calculated as the number of spikes over the duration of the sweep.

### Fast-scan cyclic voltammetry (FSCV)

Carbon fiber electrodes were made from carbon fiber (7 µm diameter, 60-90 µm length, 34-700 GoodFellow Advanced Materials) sealed within a glass pipette using horizontal puller (P-1000, Sutter Instrument). Pipettes were back-filled with potassium chloride (1 M). For recordings, carbon fibers were inserted into the slice and a triangular waveform (−0.4 to 1.3 V, 400 V/s) was delivered at 10 Hz using TarHeel CV. Electrical stimulation (40 nA, 1 ms, 1 pulse) was delivered using stimulus isolator (A365, Word Precision Instruments) and a 3 MΩ glass monopolar pipette positioned near the fiber. Fibers were calibrated in a beaker of ACSF using 400 nM dopamine steps applied every 10 seconds up to 2 µM. Peak oxidation current (at +0.6 V) were plotted against dopamine concentration and the calibration curve was calculated using a linear fit. The amplitude of dopamine transients were calculated by finding the peak within 200 ms of stimulation,

### Ex vivo 2-photon imaging

Changes in GRAB-DA3m a dLight1.3b fluorescence were recorded using a resonant-galvo scanning 2-photon micrscoscope (Ultima, Bruker) and Prairie View Software (Bruker). Excitation (920 nm) was provided by a tunable, pulsed Ti-Sapphire laser (Mai Tai DeepSee, Spectra-Physics). Emitted light was collected through a 60x objective lens (N.A 1.00; Olympus), filtered (525/70m-2p), and detected using a GaAsP PMT (7422PA40, Hamamatsu). Slices were prepared and recordings were performed as described for whole-cell patch clamp recordings with the ACSF containing the following drugs (in µM): 10 DNQX, 100 picrotoxin, 0.3 CGP 35348, 1.0 DHβE, 0.3 scopolamine, and 0.5 sulpiride. For dLight1.3b recordings, 200 µM DETQ (positive allosteric modulator for the human D1R) was also included in the ACSF^62^. Electrical stimulation (40 µA, 0.7 ms) were delivered using the methods described for slice electrophysiology. Rasterized images (∼40×40 µm, pixel size 0.068 µm) were collected at ∼30 Hz. For each slice, we recorded at two separate locations. At each locations, four fields of view were recorded around the stimulating electrode. The average of these four field of views was considered a single n, resulted in two n per slice.

Experiments involving Ca^2+^ and cAMP activity were performed using a custom-built BX51WI microscope (Olympus). Excitation (920 nm) was provided by a tuneable, pulsed Ti:Sapphire laser (Chameleon Ultra I; Coherent) and scanned using a pair of X-Y galvanometer mirrors (6215; Cambridge Technology). Emitted light was collected by a 60x objective lens (N.A. 1.00; Olympus) and filtered (ET680sp and ET525/50m-2P; Chroma) before detection by GaAsP photomultiplier tube (PMT, H10770PA-40; Hamamatsu). We used a combination of imaging (Toronado; B.W. Strowbridge), electrophysiology (Axograph X, Axograph Scientific), and custom code (Arduino) for data acquisition.

Dopaminergic modulation of voltage-gated calcium channels (VGCC) was assessed using 2-photon imaging of iSPNs identified by SYFP2 expression (AAV9-AiE0452h-minBG-SYFP2). Cells were patched, filled with the Ca^2+^ sensor Fluo5F, and held in current clamp at −80 mV. Activation of VGCC by backpropagating action potentials was induced by a brief current pulse (200-400 pA, 100 ms). Current injections were alternatively paired with electrical stimulation (20 µA, 0.5 ms) to evoke endogenous dopamine release. Electrodes were filled with 10 µM SR101 and positioned adjacent to a iSPN dendrites under 2-photon guidance at 810 nm. High-speed (2 kHZ) ‘spot-photometry’ in which the laser was centered at a manually defined region of interest on the dendrite was controlled used to measure changes in fluorescence at 810 nm.

G-Flamp2 was selectively expressed in dSPNs (pAAV-AiE0779m_3xC2-minBG-iCre(R297T)-BGHpA and AAV2/9-hSyn-DIO-opt-GFlamp2) and imaged at 920 nm. Striatal dSPNs were filled with AF594 (10 µM) to identify dendritic arbors and a stimulating electrode positioned under 2-photon guidance. Broad spatial dynamics of G-Flamp ΔF/F0 were measured from dendrites of iSPNs by collecting rasterized image sequences (30 x 40 µm, at 3 Hz), while hotspot changes were measured spot photometry in continuous 2 s segments. Traces were later aligned to stimulation and down-sampled to 50 Hz.

### 2-photon Imaging Analysis

GRAB-DA3m and dLight1.3b experiments were analyzed in Matlab. For GRAB-DA3m data, rasterized images were converted to ΔF/F_0_ using the following steps: 1) smooth individual frames using a 0.95 µm square median filter, 2) average the last 10 frames of the baseline period to get F_0_, 3) then lastly, subtract F_0_ from each frame and divide the difference by F_0._ Mean ΔF/F was calculated as the average of the entire field. The average mean peak ΔF/F of responses was calculated by identifying regions whose raw intensity signal was above 75 at baseline and whose ΔF/F response was above stated threshold, then taking the average. For both the mean and mean peak responses, we selected frames around the peak response following stimulation (∼70 ms after a single pulse or ∼155 ms after the onset of a burst stimulus).

To spatially correlate ΔF/F responses across trials, frames were divided into 1µm square grids of ‘pixels.’ A corresponding shuffled image was then created by randomly repositioning each pixel. The *corr2* function in matlab was then used to linearly correlate pixels across trials for both real and shuffled data. Correlations between frames were then averaged to get a mean correlation (R).

Data acquired using dLight1.3b expressed in dopamine axons was analyzed as for GRAB-DA3m with minor changes. Images were filtered at ∼0.5 µm. The red channel was averaged across the baseline and a threshold of 2 standard deviations above the mean was used to identify dopamine axons. Changes in ΔF/F were only quantified for regions above this threshold. Mean ΔF/F was the average change within all identified regions, while mean peak value was the average value within these regions whose response was also above 2 ΔF/F.

### Monoamine Tissue Content

To measure dopamine, HVA, and DOPAC tissue content in RIM cKO and littermate control mice, animals were deeply anesthetized with isoflurane and perfused with cold HBSS. The brain was dissected into cold HBSS and sectioned into slices with a razor blade. Portions of the dorsolateral striatum were then manually dissected, frozen on dry ice, and stored at 80°C. Tissue catecholamine levels were measured by HPLC with coupled electrochemical detection at the Vanderbilt Neurochemistry Core.

### Generation and Validation of Tetanus Toxin Light Chain AAV

Tetanus toxin light chain (TeNT-LC) was cloned in the inverted configuration into a custom double floxed inverted open reading frame (DIO) human synapsin (hSyn) promoter AAV plasmid. The construct also contains halotag, located at the N-terminus, separated from TeNT-LC by a T2A sequence. Halo-T2A-TeNT AAV (DJ serotype) was generated as previously described^63^. Briefly, HEK293T cells were co-transfected with the AAV vector along with helper plasmids (pDJ and pHelper) using calcium phosphate transfection. 72 h post-transfection cells were harvested, lysed, and purified over an iodixanol gradient column (2 h at 63,500 rpm in a Beckman Type80Ti rotor). Virus was dialyzed to remove excess iodixanol and aliquoted and stored at −80°C until use.

To validate cre-dependent TeNT-LC expression and cleavage of its native substrate, VAMP2, we infected cultured hippocampal neurons (prepared as described^63^) with pAAV DIO hSyn-Halo-2A-TeNT, along with a second AAV expressing cre recombinase (or without cre recombinase AAV as a negative control) at day *in vitro* (DIV) 10. At DIV16-20, cultures were fixed and stained with a VAMP2 antibody that does not recognize VAMP2 following TeNT-cleavage (Synaptic Systems, mouse monoclonal RRID:AB_887811). Labeling and imaging were carried out as described^64^.

### Immunohistochemistry & Microscopy

Animals were anesthetized with isoflurane, then perfused with phosphate buffered saline (PBS) followed by 4% paraformaldehyde (PFA) in PBS. Dissected brain were transferred to 4% PFA overnight and switched to a 30% sucrose solution. Slices used for ex vivo recordings were directly transferred to 4% PFA and then sucrose. After sinking, brains were blocked, tissue affixed to a chuck with O.C.T, and frozen at −20°C. Sections were then cut coronally on a cryostat (LIECA CM1510) at 30 µm thickness and stored in PBS. When quantifying SPN spine density, tissue was sectioned at 45 µm to better preserve dendritic arbors. For validating viral expression and fiber placement, sections were mounted in hardset antifade mounting media with DAPI (Vectashield).

For TH staining, sections were blocked (3% normal goat serum, 0.1% Triton-X, 1x PBS) for 2 hours at room temperature. Sections were then incubated overnight at 4°C with primary antibody (1:500 rabbit α TH) in blocking buffer. After 4-5 washes in PBS, sections were incubated for 2 hours in secondary antibody (1:500 AlexFluor-488 α rabbit) for 2 hours at 4°C. Tissue was then washed 4-5 times with PBS and mounted as described above. For spaghetti monster staining, sections were blocked for 2 hours and the fluorescently labeled primary antibody (1:100 AlexaFluor-488 α myc) was applied in blocking buffer overnight at 4°C. Sections were then washed and mounted as described above.

To validate viral expression, fiber placement, and TH knockout, slides were imaged on a slide scanning microscope (Olympus) at 10x or 20x magnification. For quantifying SPN spine density, slides were imaged on spinning disk confocal (3i) at 63x.

Quantification of striatal TH for experiments involving TH knockout was performed in ImageJ (Fiji) using prepacked tools. In sections ipsilateral and contralateral to the injected hemisphere, square regions of interest were drawn in the dorsal striatum and the corpus collosum. The mean intensity of the corpus collosum was then subtracted from that of the dorsal striatum on each side to account for background fluorescence. The reduction in TH was then calculated as average fluorescent intensity of the ipsilateral side divided by the contralateral side.

Spine density was manually measured in ImageJ using prepackaged tools. A 20 µm and 40 µm radius was drawn from the center of the soma to demarcate proximal and distal dendrites. Dendrites were then traced along their length and spine heads marked manually. Spine density was calculated as the number of spines divided by the length of dendrite.

### Fiber photometry

Following 3 weeks of recovery in their home cage, mice were habituated for a minimum of 5 days to locomote while head-fixed on a cylindrical wheel placed into a dark soundproof chamber and were water restricted to incite consumption of water rewards from a spout at semi-random 10 to 30 s intervals. Licking was monitored by a capacitive touch sensor (AT42QT1010; Sparkfun). Extracellular DA levels in striatum were imaged by providing constant 470 nm excitation light from a LED (M470F3, Thorlabs; 30 uW measured at tip of patch cord) coupled to a fluorescence mini-cube (FMC3_E(460–490)_F(500–550)_S; Doric) and connected to the mouse via a fiber optic patch cord (400 μm, 0.48 NA; Doric) and zirconia sleeve. Emission light was collected through the same patch cord and mini-cube by a femtowatt photoreceiver (2151; Newport) fitted with a FC adapter (FOA_2151_FC; Doric) via a 600 um 0.48NA fiber optic (Doric), continuously digitized at 2 kHz using a National Instruments acquisition board (USB-6343) and Wavesurfer software (Janelia).

For analysis of fiber photometry data, photometry traces were lowpass filtered at 50 Hz, down-sampled to 50 Hz, and smoothed by a sliding 5 ms average window. Raw signals were converted to ΔF/F by dividing the traces by the mean value of the entire recording. Spontaneous transients were identified by finding the difference between local minima and local maxima. A minimum peak threshold of 1.5% ΔF/F was used as inclusion criteria, as peaks larger than this threshold were rarely detected in mice expressing GFP or following administration of the D1R antagonist SKF-38393. Mean transient amplitude was calculated by averaging the amplitude of all identified transients within a session. Transient rate was calculated as the number of transients divided by duration of the included recording. To exclude reward related transients, transients were excluded if they occurred within 2 seconds of reward delivery.

For quantification of lick-rate data, capacitor activation events were considered licks if they were between 5-50 ms in duration and occurred at least 5 ms apart. Licks were then counted as the remaining number of sensor onsets events.

### Drug Administration

Reserpine was dissolved in the following solution (10% DMSO, 40% polyethylene glycol 400, 5% Tween80, 45% normal saline) at a concentration of 0.1 mg/mL. The same solution without reserpine was used as vehicle. Mice received an i.p. injection of reserpine (0.5 mg/kg) or vehicle the night prior to slice or behavioral experiments. For the accelerating rotarod assay, doses were spaced 48 hours apart to allow time for drug washout in between doses.

### Open Field

All behavioral experiments were performed during the light period. Prior to the open field, animals were habituated to the room for 15-20 minutes. Animals were placed in a square open field arena (50 x 50 cm) dimly illuminated by an overhead light in a dark, quiet room. An overhead IR webcam (ELP-USBFHD05MT-KL36IR) and custom Matlab code (2018b Mathworks) were used to capture video (20-30 Hz) for 30 minutes. All behavioral apparatuses were cleaned with 70% ethanol solution between animals and allowed time to aerate.

We used custom Matlab (2019a, Mathworks) code to track to track an animal’s center point from videos. Velocity traces were calculated by dividing the change in distance (cm) between frames by the change in time (s). Mean velocity for the session was the average of the velocity trace.

Percent time mobile was defined as the number of frames the animal’s velocity was greater than 2 cm/s divided by the total number of successfully tracked frames. To calculate mean peak velocity, all local maxima of the velocity trace were identified and then the average value of peaks greater than 2 cm/s was taken.

### Accelerating Rotarod

Mice were initially placed on a rotarod (Ugo Basile srl) at rest and allowed to habituate for 2-3 minutes. Animals generally performed either three trials per day for three consecutive days or five trials per day for five consecutive days. For an individual trial, the rotarod wheel started at 5 rpms and accelerated linearly to 80 rpms over the course of 300 seconds. We quantified the latency to fall, which was defined as the point at which an animal failed to maintain at least a 90 degree climbing position relative to the ground (animals could not hang on the rod and were not required to physically fall off). Trials were separated by a 3 minute rest period during which mice were returned to the rotarod wheel.

### Conditioned Place Preference

Following surgery, mice were allowed to recover for a minimum of 8 weeks. One week prior to experimentation, animals were habituated to the behavioral room and handling daily for at least an hour. Afterwards, mice were habituated to optical fiber tethering (200 µm diameter, 0.39 NA, ThorLabs) in a clean empty cage for two 30 min periods across two days. Optical fibers were connected to an LED (LEDD1B diver with M455F3, ThorLabs) through an optical commutator (RJ2, Thorlabs) to allow free movement.

For the two day baseline period, mice were allowed to freely explore a custom-built two chamber box under dim illumination for 20 minutes each day. The two chambers differed in both the patterning of the walls (vertical vs. horizontal stripes) and the texture of the floor (holes vs. slates). No light was delivered during the baseline period. Individual baseline preference (percent of total time) for each chamber was calculated and used to assign the light paired chamber for each animal. Assignment was made so that 1) the average preference within group was as close to 50% as possible and 2) the two chambers were equally represented within each group. Video recordings, real-time positional tracking, and light delivery were controlled with custom Matlab (Mathworks) code.

On the subsequent two days, mice underwent the 30 min real-time place preference test (RTPP) sessions. During RTPP, regular bursts of blue light (455 nm, ∼2mW, 10 pulses, 5 ms duration, 20 Hz) were delivered every 8 seconds within the light-paired chamber. The task was designed so that mice could not trigger more frequent stimulation by crossing back and forth. Preference was calculated as the percent of total time spent in the light-paired chamber.

After RTTP, mice underwent one conditioned place preference (CPP) test session the following day for 30 minutes. Conditions were maintained as for the RTPP sessions, but without light delivery.

### Optogenetic Stimulation in the Open Field

Optogenetic stimulation in the open field was performed after completion of conditioned place preference testing. Mice bilaterally tethered were placed in a square open field arena (50 x 50cm) in the dark. Mice were allowed to locomote freely for 30 minutes, during which optical stimulation (455 nm, ∼2mW, 10 pulses, 5 ms duration, 20 Hz) was delivered at random intervals (5-20 s). Animals were filmed using an infrared camera capturing frames at ∼20 Hz. Centerpoint tracking and calculation of continuous velocity were performed using custom Matlab code. To remove high amplitude artifacts, velocities >40 cm/s were excluded, and velocity traces were then smoothed using a slide average of 0.75 s. Mice were considered immobile prior to stimulation if they did not exceed a velocity of 5 cm/s for 3.5 s prior to stimulation onset. Velocity traces were then normalized to mean value between −1 and −0.5 s. Change in velocity was measured at 0.75 to 1.25 s post stimulation.

### Statistics

Statistical comparisons were planned in advance and only performed once upon completion of data acquisition. We did not perform power calculations, but estimated group sizes based on prior experiments from the within the lab and from the literature. No assumptions were made about the normality and we did not test data for normality. Wilcoxon rank-sum or signed-rank tests were used for comparisons across two groups. Kruskal-Wallis tests followed by Dunnet’s or Tukey’s post hoc tests were used for comparisons across three or more groups. All statistical comparisons were performed using Matlab.

### Data and code availability

The data, code, protocols, and key lab materials used and generated in this study are listed in a Key Resource Table alongside their persistent identifiers.

## Acknowledgements

We thank James Taniguchi and Akul Prakash for helping acquiring photometry data. We thank Tracy Frost for managing the animal colony.

## Funding

This work was funded in part by Aligning Science Across Parkinson’s (ASAP-020529) through the Michael J. Fox Foundation for Parkinson’s Research (MJFF) (CPF), Aligning Science Across Parkinson’s Discovery Fellowship (ASAP 028263) (MMM), R01MH134957 (MJK), R35NS116879 (MJK), funding from a McGill Conrad F. Harrington Fellowship and CIHR Postdoctoral Research Award (SP), R21NS133640 (NXT), CIHR (551684) (NXT) and the Canada First Research Excellence Fund ‘Healthy Brains, Healthy Lives’ at McGill University (NXT).

For the purpose of open access, the author has applied a CC BY public copyright license to all Author Accepted Manuscripts arising from this submission.

## Author contributions

MMM, NXT and CPF conceived and designed the project. MMM and CPF wrote the manuscript with input from other authors. AGY performed and analyzed data for 2P imaging of endogenous receptor signaling, SE performed and analyzed data for dose response measurements and assisted with data acquisition, JA performed and analyzed data for iSPN enhancer viral targeting, SKP, RM and NXT performed and analyzed data for fiber photometry experiments, CSW and MJK designed and performed validation experiments using tetanus toxin construct. MMM did all other experiments and analysis, interpretation of data and preparation of figures.

## Competing interests

The authors declare no competing interests.

---

**Supplemental Figure 1:**
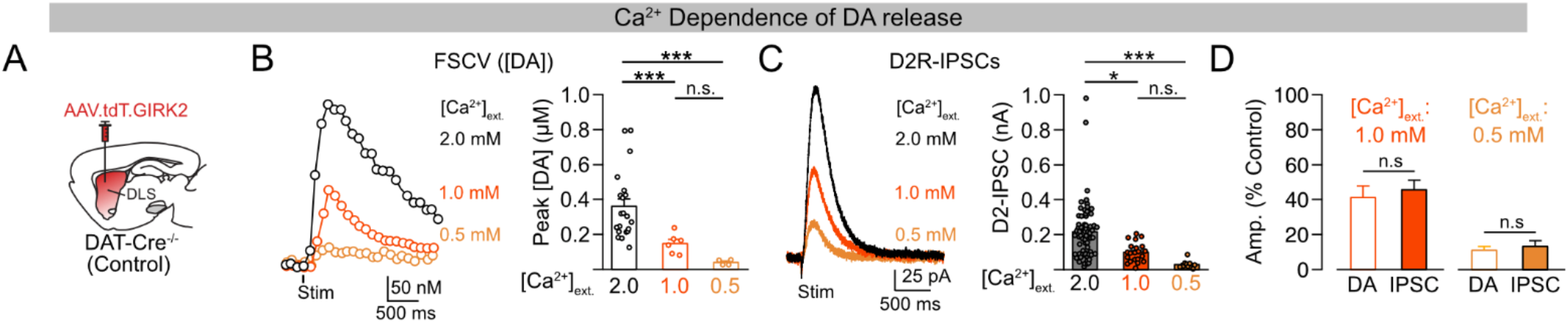
Lowering extracellular [Ca^2+^] equally reduces the amplitude of dopamine transients and D2R-IPSCs. A. Schematic of approach to measure dopamine transients (FSCV) and D2R activation (D2R-IPSCs) by virally expressing GIRK2 channels in iSPNs of control animals. B. Examples traces and summary of dopamine transient amplitudes in 2.0, 1.0, and 0.5 mM extracellular Ca^2+^. Kruskal-Wallis test (*** p<0.001) followed by Tukey’s test (2 mM vs. 1 mM: *** p<0.001, 2mM vs 1 mM: *** p<0.001. 1 mM vs 0.5 mM: n.s. p>0.05). C. Examples traces and summary of D2R-IPSC amplitudes in 2.0, 1.0, and 0.5 mM extracellular Ca^2+^. Kruskal-Wallis test followed by Tukey’s test (*** p<0.001) followed by Tukey’s test (2 mM vs. 1 mM * p<0.05, 2mM vs 1 mM *** p<0.001. 1 mM vs 0.5 mM n.s. p>0.05). D. Quantification of the relative change in D2R-IPSC and dopamine transient amplitudea for 1.0 mM (n.s. p>0.05) and 0.5 mM (n.s. p>0.05) extracellular Ca^2+^. Wilcoxon rank-sum test. Summary data are the mean ± SEM.

**Supplemental Figure 2:**
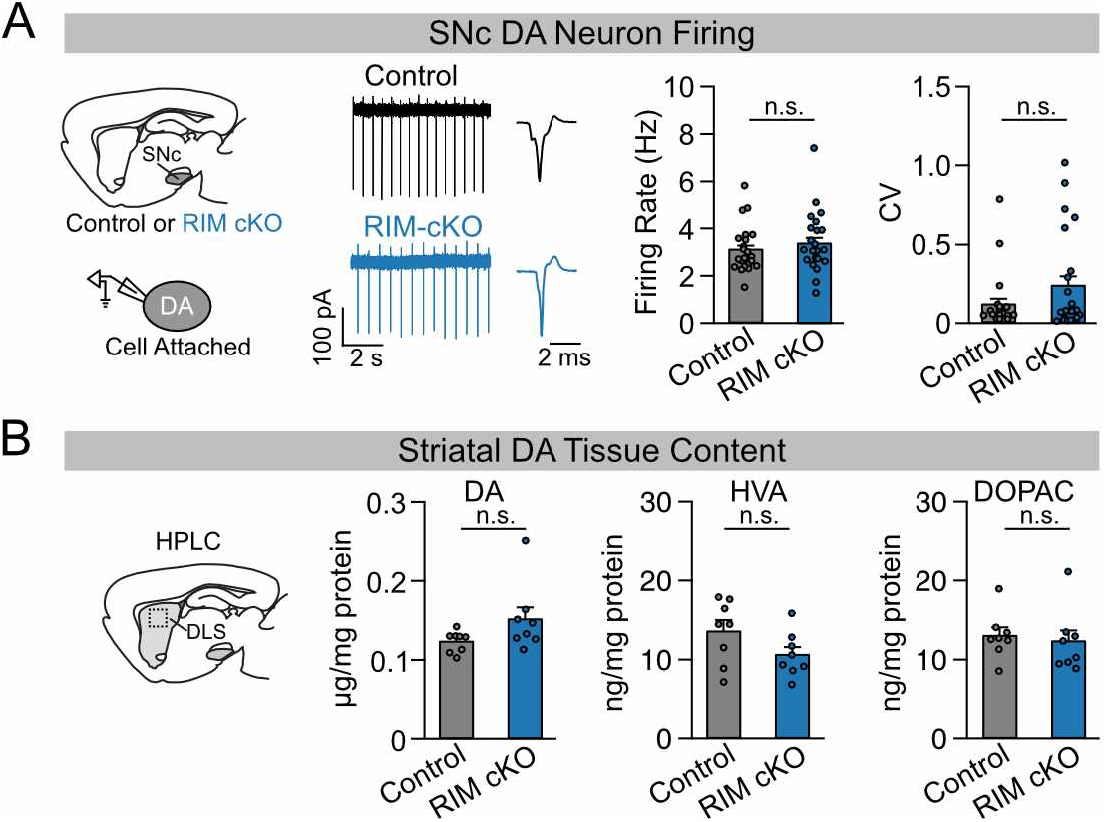
Conditional knockout of RIM1/2 does not alter dopamine neuron pacemaking activity or striatal monoamine tissue content. A. Schematic of cell-attached recording setup and examples of pacemaking activity in SNc dopamine neurons from control and RIM cKO mice (left). Quantification of intrinsic firing rate (n.s. p>0.05) and coefficient of variance in interspike internals (n.s. p>0.5) for control and RIM cKO mice (right). Wilcoxon rank-sum test. B. Summary of dopamine (DA, n.s. p>0.5), homovanillic acid (HVA, n.s. p.0.5), and 3,4-dihydroxyphenylacetic acid (DOPAC, n.s. p>0.5) tissue content in the DLS of control and RIM cKO mice. Wilcoxon rank-sum test. Summary data are the mean ± SEM.

**Supplemental Figure 3:**
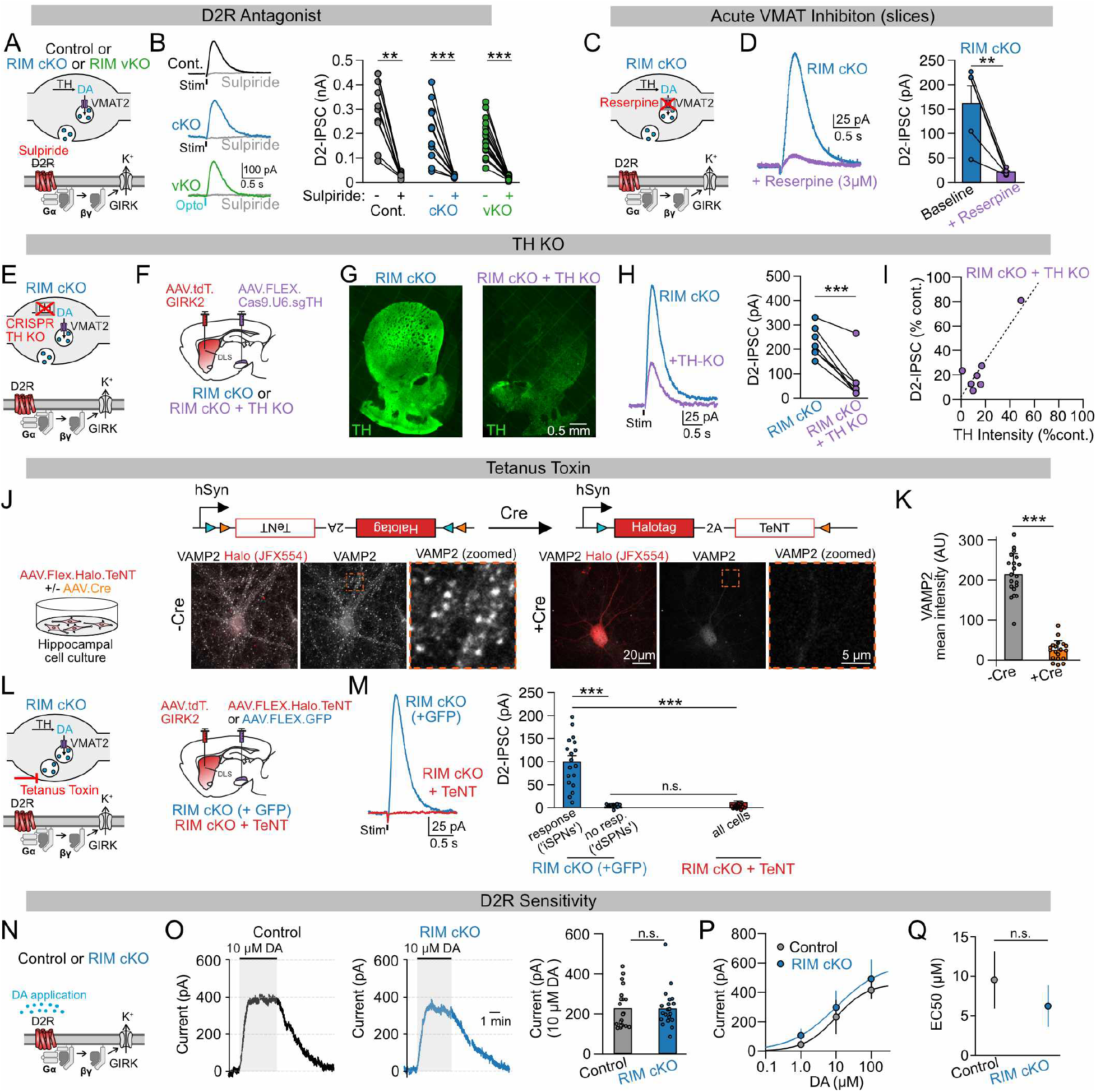
Vesicular dopamine release from dopamine neurons drives D2R-IPSCs in RIM cKO mice without an appreciable upregulation D2R-mediated signaling. A. Schematic depicting the blockade of D2Rs with the antagonist sulpiride in slices from control (Cont.), RIM cKO (cKO), or RIM vKO (cKO) mice. B. Example traces and quantification of sulpiride’s effect on D2R-IPSCs in control (** p<0.01), RIM cKO (*** p<0.001), and RIM vKO mice (*** p<0.001). Wilcoxon sign-rank. C. Schematic depicting the blockade of the vesicular monoamine transporter 2 (VMAT2) with reserpine in slices from RIM cKO mice. D. Example traces and quantification of reserpine’s effect on D2R-IPSCs in RIM cKO mice (*** p<0.001). Wilcoxon rank-sum test. E. Schematic depicting CRISPR-mediated knockout of tyrosine hydroxylase (TH) in dopamine neurons. F. Schematic of approach to measure D2R-IPSCs in RIM cKO mice with CRISPR-mediated TH knockout. G. Example images of TH staining in the striatum of RIM cKO mice with and without CRISPR-mediated knockout of TH. H. Examples of D2R-IPSCs in iSPNs from RIM cKO mice without (RIM cKO) and with CRISPR-mediated TH knockout (TH-KO, left). Summary of TH knockout’s effect on D2R-IPSC amplitude (*** p <0.001, right). Wilcoxon sign rank test. I. Summary of averaged normalized D2-IPSC amplitudes as a function of normalized TH staining intensity in RIM cKO mice following CRISPR-mediated TH knockout. J. Schematic depicting Cre-dependent expression of tetanus toxin in a culture of hippocampal neurons (left). Example images of VAMP2 staining in neuronal cultures infected with virus encoding Cre-dependent tetanus toxin alone (-Cre) or coinfected with virus encoding Cre-recombinase (right, +Cre). K. Quantification of VAMP2 staining in cultures transfected with Cre-dependent tetanus toxin alone (-Cre) or with Cre-recombinase (+Cre, ***p<0.001). Two-sided student’s t-test. L. Schematic depicting the blockade of vesicular fusion in dopamine neurons by tetanus toxin (left). Schematic showing the approach to measure D2-IPSCs in iSPNs from RIM cKO mice expressing GFP (+ GFP) or tetanus toxin (+ TeNT) in dopamine neurons. M. Example traces of D2R-IPSCs from RIM cKO mice selectively expressing green fluorescent protein (GFP) or tetanus toxin (TeNT) in dopamine neurons (left). Quantification of D2R-IPSC amplitudes from RIM cKO mice expressing GFP (responsive and non-responsive cells) or tetanus toxin (all cells, right). Kruskal-Wallis test (*** p<0.001) followed by Tukey’s post hoc test (response vs no response, *** p<0.001, response vs. TeNT, *** p<0.001, no response vs. TeNT, n.s. p>0.05). N. Schematic of D2R-mediated GIRK currents evoked by wash-on of exogenous dopamine. O. Example of D2R-mediate GIRK2 currents in response to 10 µM dopamine application in control and RIM cKO mice (left). Quantification of the average current amplitude for control and RIM cKO iSPNs (right, n.s. p>0.05). P. Summary of D2R-mediated GIRK2 current amplitudes induced by exogenous application of 1, 10, or 100 µM dopamine in iSPNs from control and RIM cKO mice. Q. Quantification of EC50 values from (P) (n.s. p>0.5). Summary data are the mean ± SEM.

**Supplemental Figure 4:**
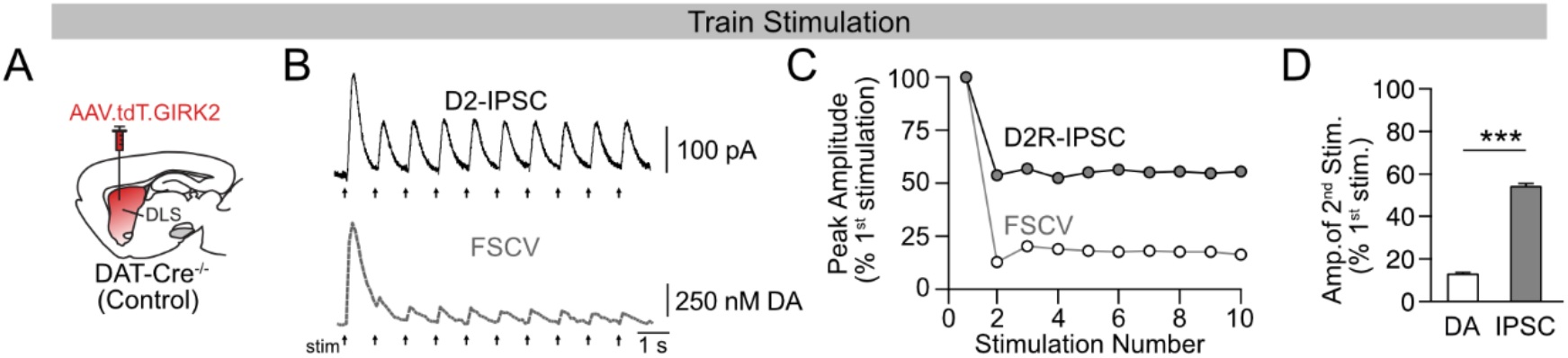
Stimulus trains reveal a disconnect between spatially diffuse dopamine release and D2R activation in wild type animals. A. Schematic of approach to measure dopamine transients (FSCV) and D2R activation (D2R-IPSCs) by virally expressing GIRK2 channels in iSPNs of control (wild type) animals. B. Examples of D2R-IPSCs and dopamine transients evoked by a 1 Hz stimulus train. C. Summary of D2R-IPSC and dopamine transient amplitudes normalized to the amplitude of the first response. D. Quantification of the relative change in amplitude of dopamine transients (DA) and D2R-IPSCs (IPSC) from the first to second pulse (*** p <0.001). Wilcoxon rank-sum test. Summary data are the mean ± SEM.

**Supplemental Figure 5:**
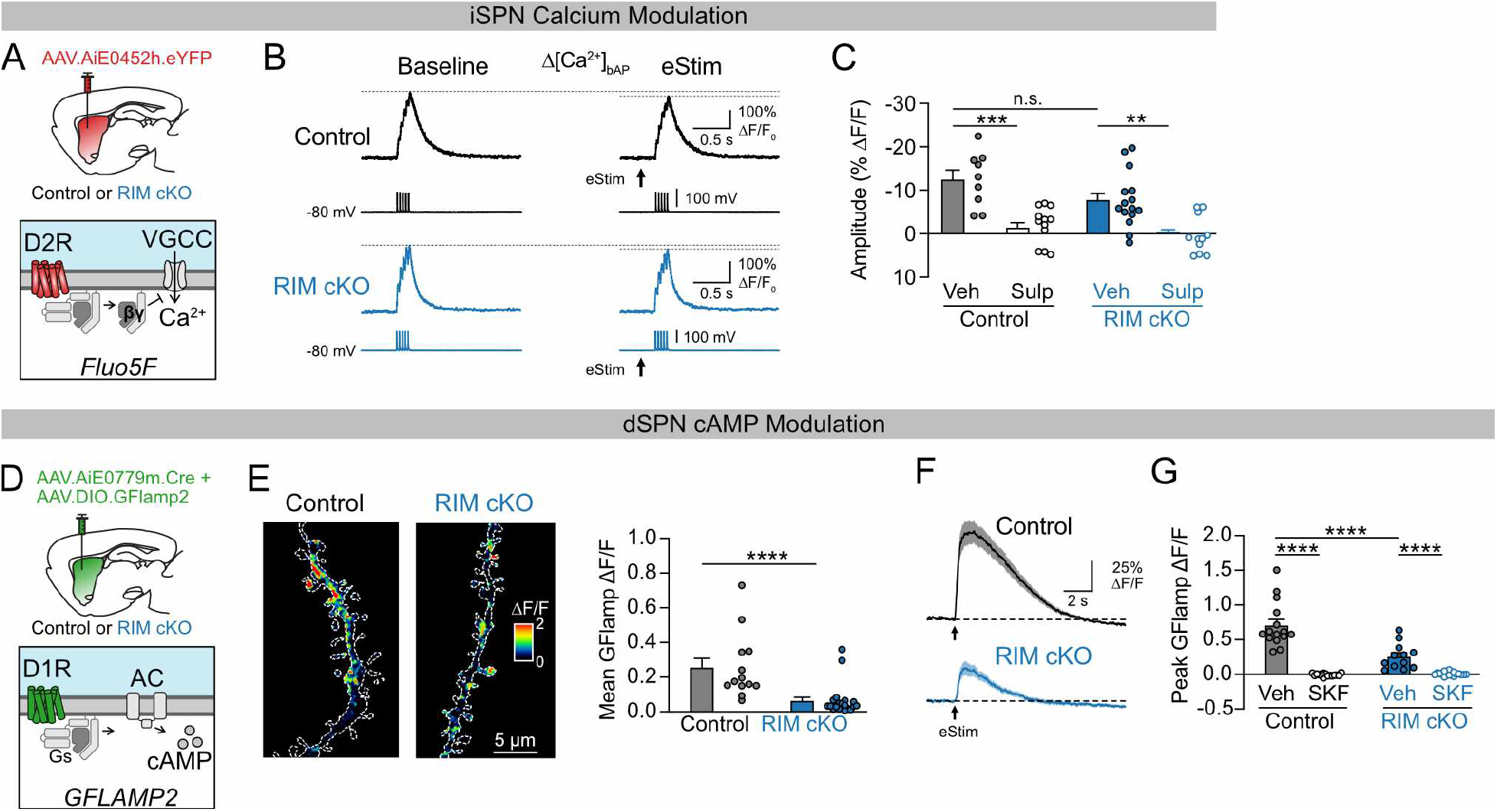
Dopamine release in RIM cKO mice engages endogenous effectors downstream of D2Rs and D1Rs. A. Schematic of experimental approach to measure D2R-mediated modulation of dendritic calcium influx induced by backpropagating action potentials in iSPNs from control and RIM cKO mice. B. Examples of D2R-mediated reductions in dendritic calcium influx. C. Quantification of D2R-mediated modulation of dendritic calcium influx in iSPNs from control and RIM cKO mice in the absence and presence of the D2R antagonist sulpiride. Kruskal-Wallis test (*** p<0.001) followed by Dunnet’s post hoc test (Control Veh vs. Control Sulp: ** p<0.001, RIM cKO Veh vs. RIM cKO Sulp: ** p<0.01, Control Veh vs. RIM cKO Veh: ns p>0.05). (Vehicle data from Fig 1H.) D. Schematic showing the experimental approach to measure D1R-mediated changes in dendritic cAMP dSPNs from control and RIM cKO mice with the fluorescent sensor G-Flamp2. E. Example of GFlamp-2 activity in response to electrical stimulation in dSPNs from control and RIM cKO mice and quantification of the average change in GFlamp-2 fluorescence (ΔF/F) across dendrites in dSPNs from control and RIM cKO mice. Wilcoxon rank-sum test. F. Example traces of GFlamp-2 activity from hotspots detected in dendrites of dSPNs from control and RIM cKO mice. G. Quantification of hotspot amplitudes in dSPNs from control and RIM cKO mice in the absence and presence of the D1R antagonist SKF38393. Kruskal-Wallis test (*** p<0.001) followed by Dunnet’s post hoc test (Control Veh vs. Control SKF: *** p<0.001, RIM cKO Veh vs. RIM cKO SKF: ** p<0.01, Control Veh vs. RIM cKO Veh: ns p>0.05). Summary data are the mean ± SEM.

**Supplemental Figure 6:**
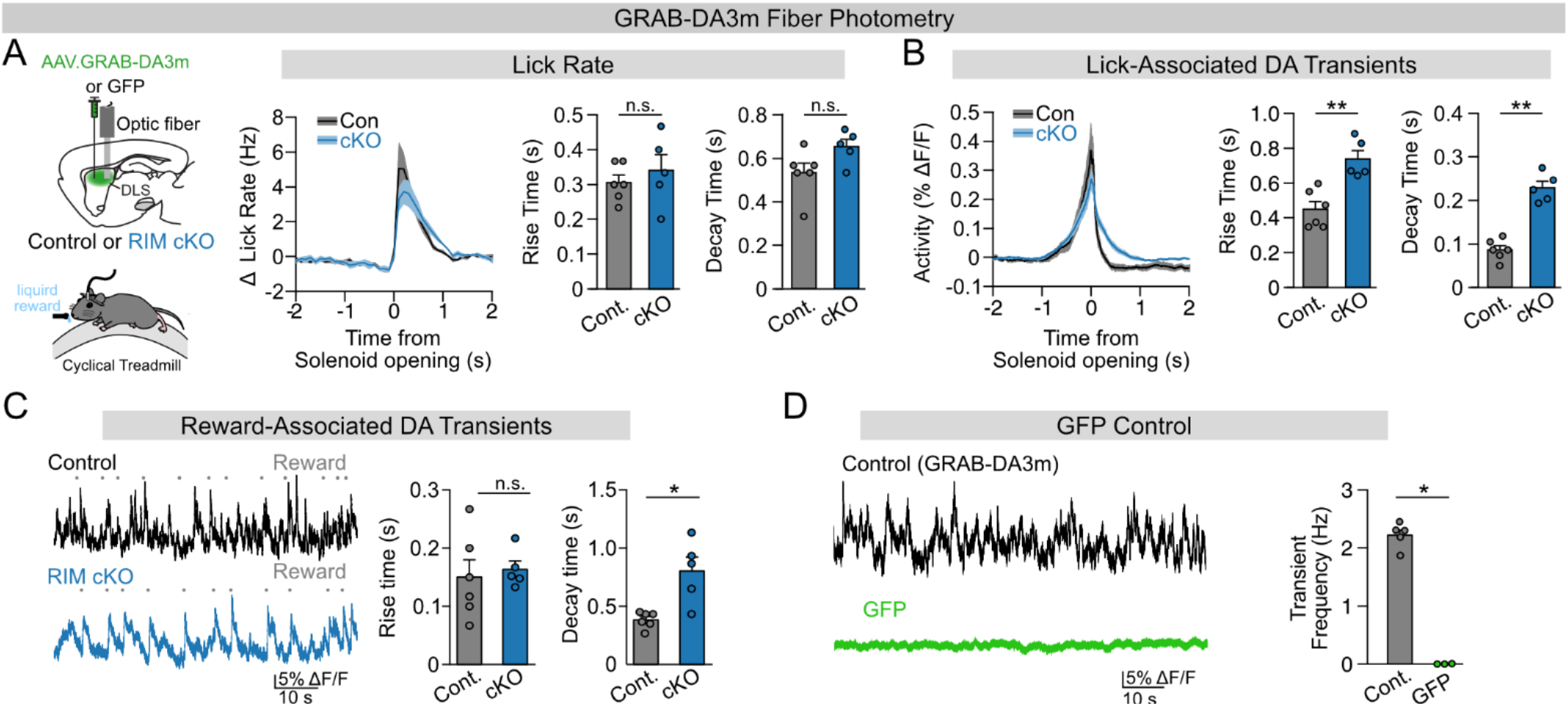
RIM cKO mice show subtle effects on reward-related measures of behavior and dopamine activity. A. Schematic depicting the approach to measure bulk dopamine release with the fluorescent sensor GRAB-DA3m in mice periodically receiving a liquid reward (left). Summary of lick rate around reward delivery (solenoid opening) in control (Cont.) and RIM cKO (cKO) mice (middle). Quantification of the time to peak (ns p>0.05) and decay time (ns p>0.05) for licking data in the middle panel (right). Wilcoxon rank-sum tests. B. Summary of GRAB-DA3m activity (ΔF/F) aligned to licks for control and RIM cKO mice (left). Quantification of time to peak (** p<0.01) and decay time (** p<0.01) for the GRAB-DA3m data shown to the left (right). Wilcoxon rank-sum tests. C. Example of GRAB-DA3m activity during reward delivery (grey dots) from control and RIM cKO mice (left). Quantification of rise times (n.s. p>0.05) and decay times (* p<0.05) for reward evoked transients in control (Cont.) and RIM cKO mice (cKO). Wilcoxon rank-sum tests. D. Example photometry traces from control animals expressing GRAB-DA3m or green fluorescent protein (GFP, left). Quantification of transient frequency in mice expressing GRAB-DA3m (Cont.) and GFP (** p<0.01). Wilcoxon rank-sum test. Summary data are the mean ± SEM.

**Supplemental Figure 7:**
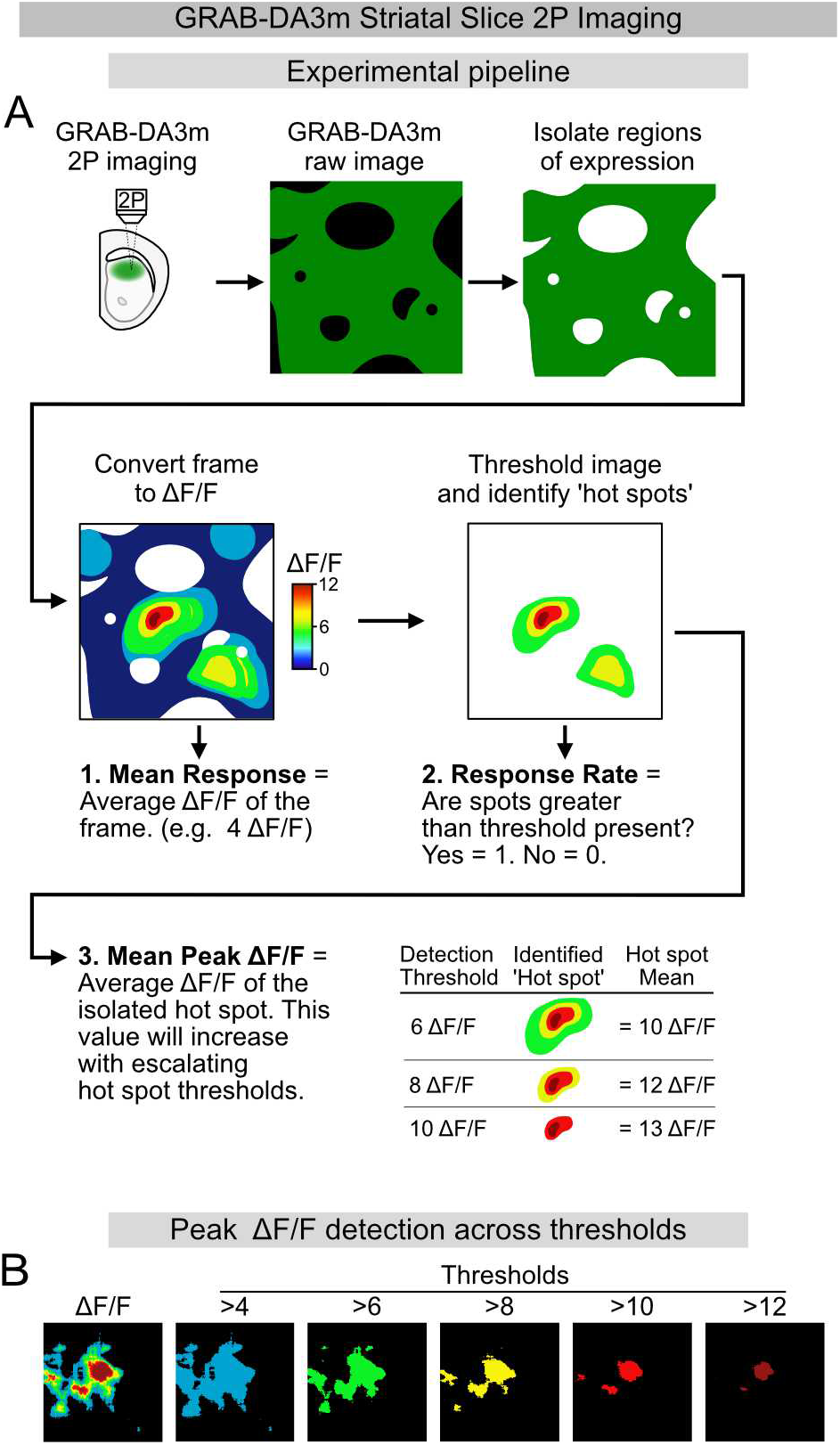
Method for quantifying measures of dopamine release hotspots. A. Schematic depicting the pipeline for detecting and analyzing hotspots of dopamine release. B. Example of a dopamine release hotspot across a range of ΔF/F thresholds.

**Supplemental Figure 8:**
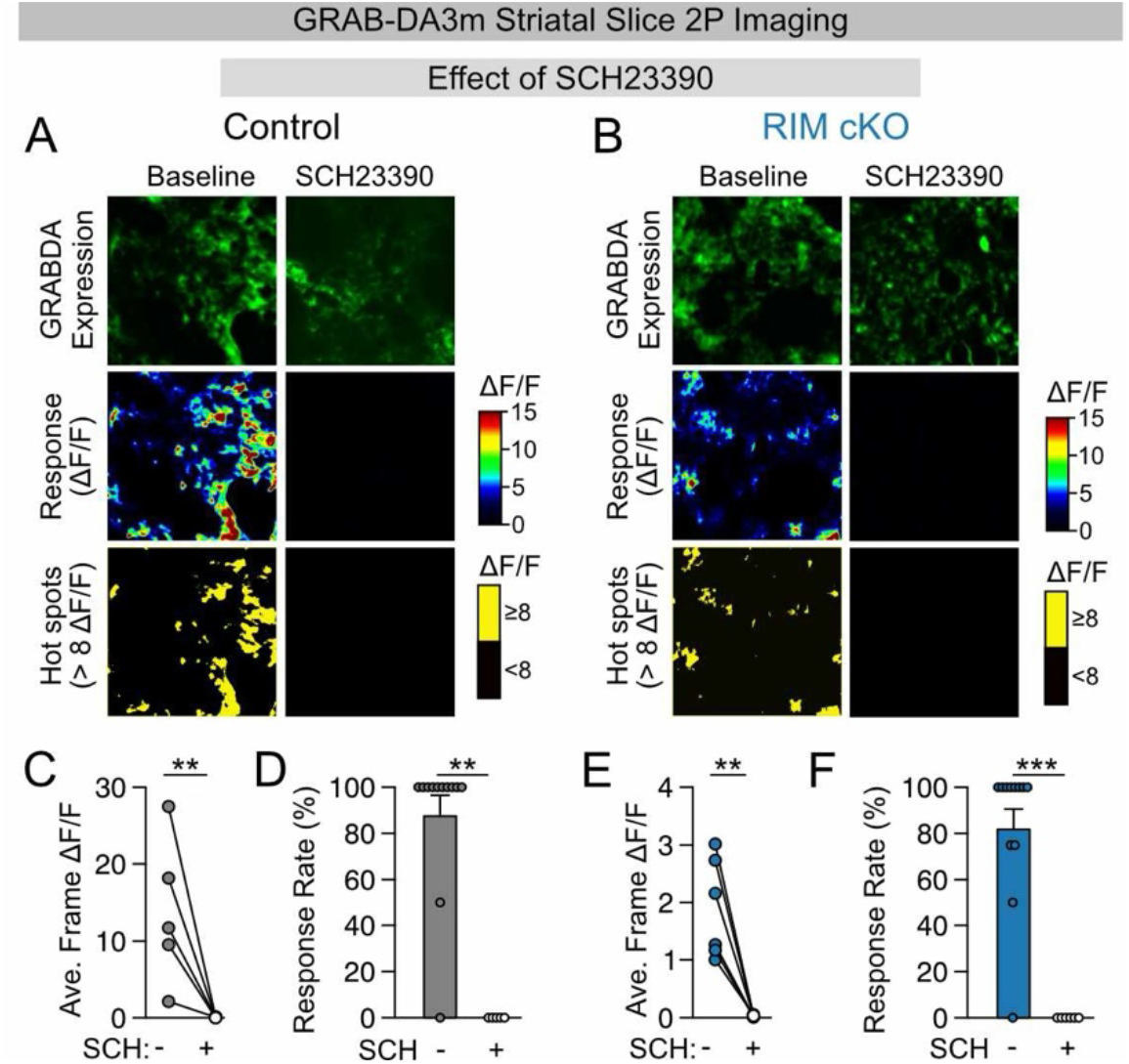
The D1R antagonist SCH23390 blocks electrically evoked increase in GRAB-DA3m fluorescence and eliminates dopamine hotspots in both control and RIM cKO mice. A. Examples of GRAB-DA3m expression, electrically evoked GRAB-DA3m activity (ΔF/F), and isolated hotspots (regions > 8 ΔF/F) in a control animal in the absence and presence of the D1R antagonist SCH23390. B. Similar experimental approach as shown in (A), but from a RIM cKO mouse. C. Quantification of the average GRAB-DA3m response (ΔF/F) from control mice in the absence and presence of SCH23390 (** p<0.01). Wilcoxon sign-rank test. D. Quantification of response rate (presence of hotspots > 8 ΔF/F) in control mice in the absence and presence of SCH23390 (*** p<0.001). Wilcoxon rank-sum test. E. Quantification of the average GRAB-DA3m response (ΔF/F) for slices from RIM cKO mice in the absence and presence of SCH23390 (** p<0.01). Wilcoxon sign-rank test. F. Quantification of response rate (presence of hotspots > 8 ΔF/F) in RIM cKO mice in the absence and presence of the SCH23390 (*** p<0.001). Wilcoxon rank-sum test. Summary data are the mean ± SEM.

**Supplemental Figure 9:**
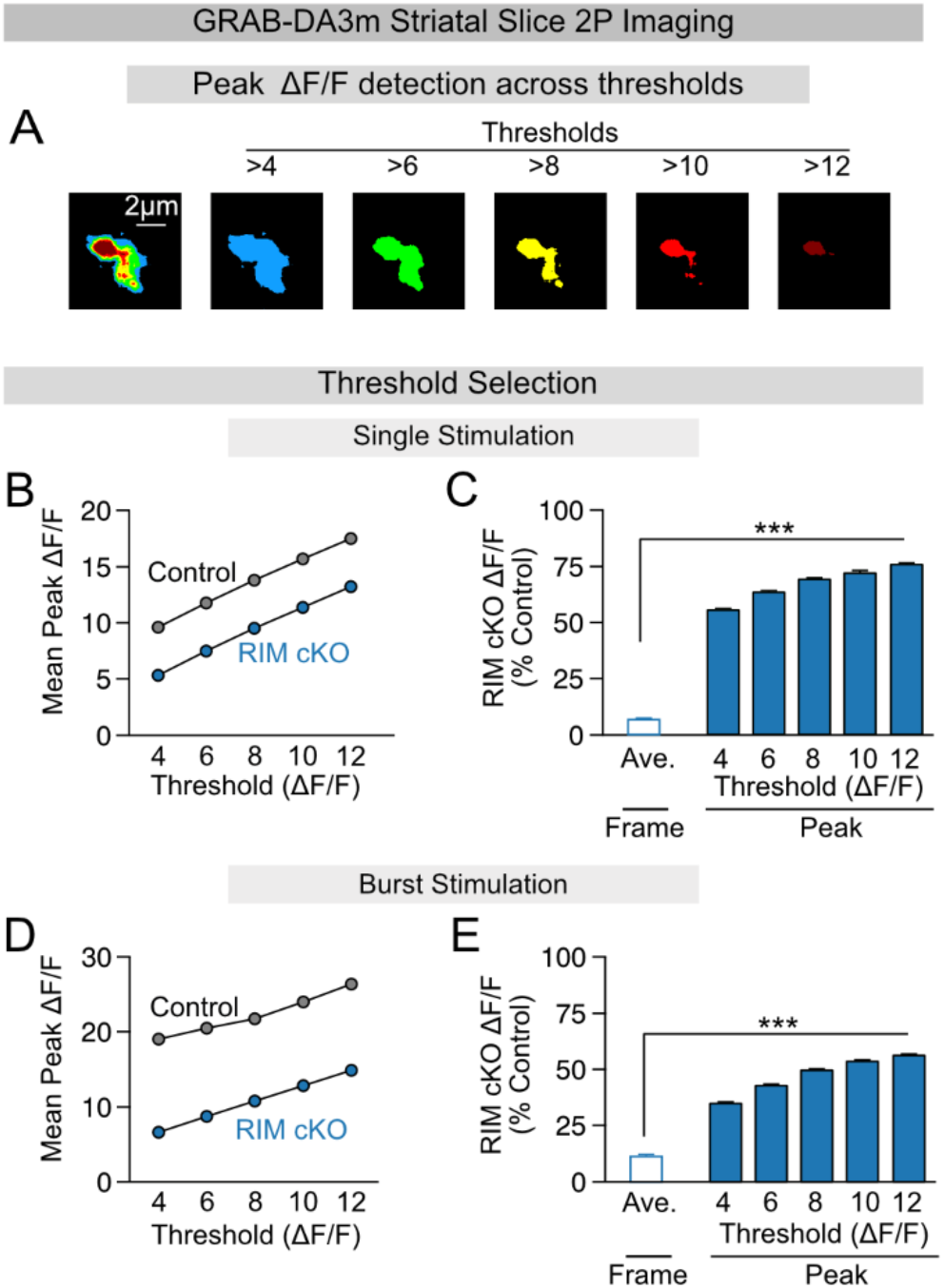
The differential effect of RIM1/2 knockout on mean response and mean peak response persists across a range of ΔF/F thresholds. A. Example of a dopamine hotspot across a range of ΔF/F thresholds. B. Summary of the average mean amplitude within dopamine hotspots (mean peak) in response to a single electrical stimulus across a range of detection thresholds for control and RIM cKO mice. C. Quantification of the relative reduction in mean and mean peak amplitudes across a range of detection thresholds for RIM cKO mice compared to controls. Data is for a single electrical stimulus. Kruskal-Wallis test (*** p<0.001) followed by Tukey’s post hoc test (Ave. vs 4, 6, 8, 10, 12: *** p<0.001). D. Summary of mean peak amplitudes in response to a burst stimulus (40 Hz, 5 pulse) across a range of detection thresholds for control and RIM cKO mice. E. Quantification of the relative reduction in mean and mean peak amplitudes across a range of detection thresholds for RIM cKO mice compared to controls. Data is for a burst stimulus as described in (D). Kruskal-Wallis test (*** p<0.001) followed by Tukey’s post hoc test (Ave. vs 4, 6, 8, 10, 12: *** p<0.001). Summary data are the mean ± SEM.

**Supplemental Figure 10:**
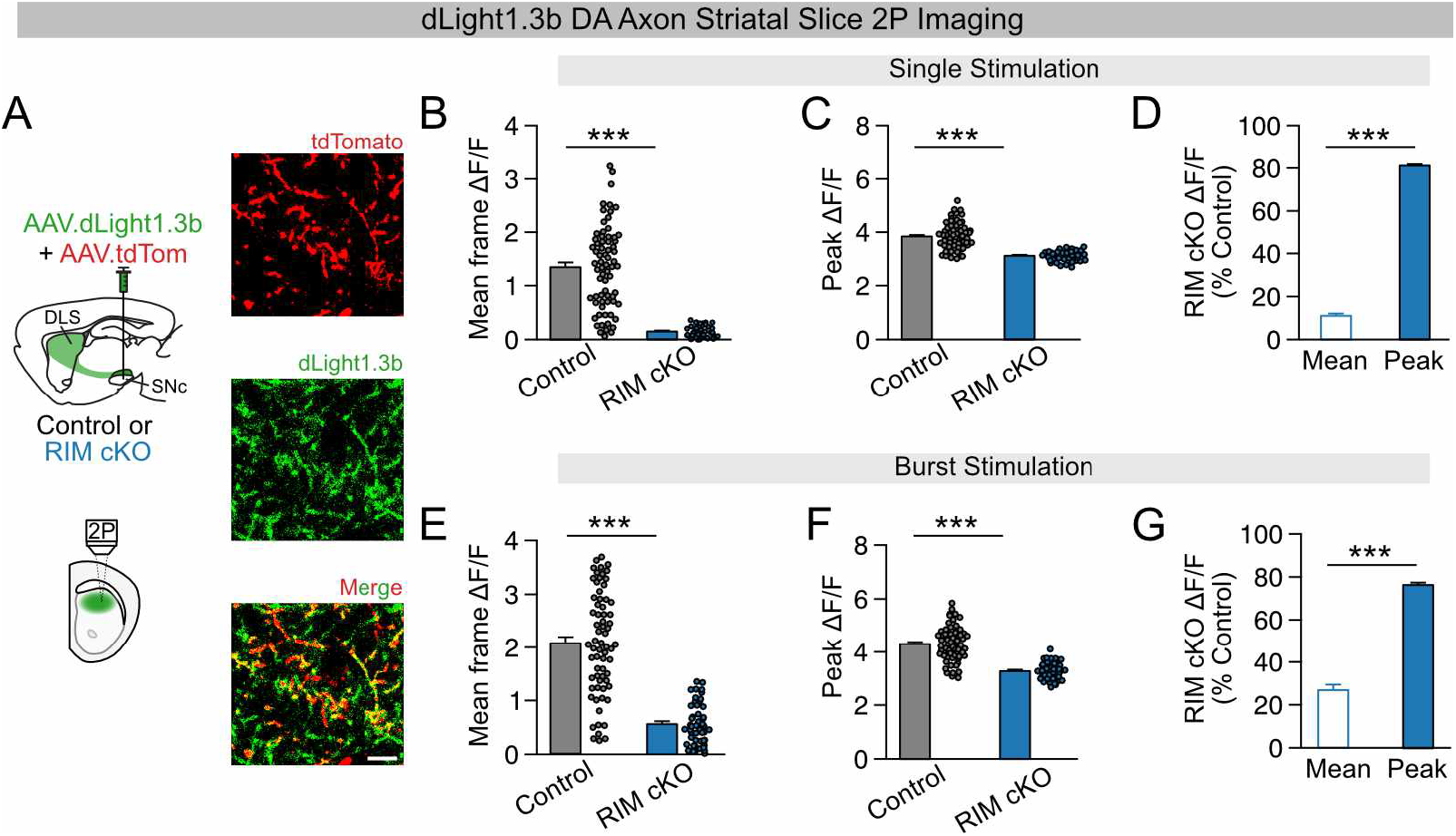
Hotspots of dopamine release in RIM cKO mice are also detectable with the fluorescent dopamine sensor dLight1.3b. A. Schematic depicting the approach to measure dopamine release in slices from mice expressing dLight1.3b and tdTomato in nigrostriatal dopamine axons (left). Example images of dopamine axons within the dorsolateral striatum labeled with dLight1.3b and tdTomato (right). B. Quantification of the average dLight1.3b response (ΔF/F) for a single electrical stimulus in slices from control and RIM cKO mice (*** p<0.001). Wilcoxon rank-sum test. C. Quantification of the average mean amplitude within dopamine hotspots (ΔF/F > 2.0) evoked by a single electrical stimulus for control and RIM cKO mice (*** p<0.001). Wilcoxon-rank sum test. D. Relative decreases in average dLight1.3b response and average mean hotspot amplitude evoked by a single stimulus for RIM cKO mice compared to controls (*** p<0.001). Wilcoxon rank-sum test. E. Quantification of the average dLight1.3b response (ΔF/F) to a burst stimulus (40 Hz, 5 pulses) in slices from control and RIM cKO mice (*** p<0.001). Wilcoxon rank-sum test. F. Quantification of the average mean amplitude within dopamine hotspot (ΔF/F > 2.0) evoked by a burst stimulus for control and RIM cKO mice (*** p<0.001). Wilcoxon rank-sum test. G. Relative decreases in average dLight1.3b response and average mean hotspot amplitude evoked by a burst stimulus for RIM cKO mice compared to controls (*** p<0.001). Wilcoxon rank-sum test. Summary data are the mean ± SEM.

**Supplemental Figure 11:**
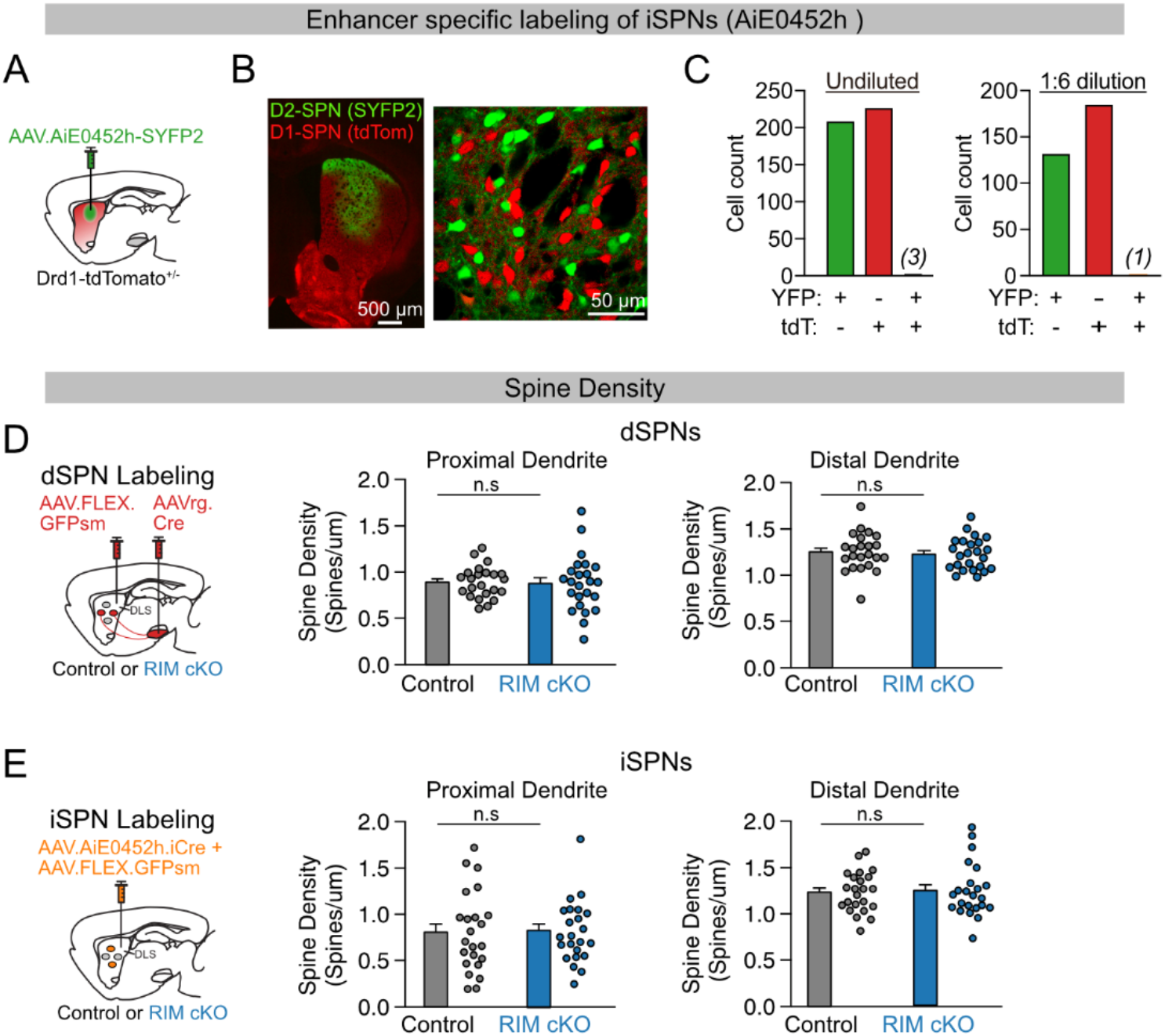
Validation of viral strategy for isolating iSPNs and breakdown of SPN spine density in proximal and distal dendrites. A. Schematic depicting the approach to label iSPNs with eYFP using an iSPN specific enhancer AAV. B. Example image of eYFP expression in the striatum of a D1-tdTomato mouse. C. Counts of eYFP+, tdTomato+ (tdT), and double labeled cells in mice injected with undiluted or diluted (1/6) virus. D. Schematic showing the experimental approach for quantifying spine density in dSPNs (left). Quantification of spine density in proximal (n.s. p>0.5) and distal dendrites (n.s. p>0.5) of dSPNs from control and RIM cKO mice (right). Wilcoxon rank-sum test. E. Schematic showing the experimental approach for quantifying spine density in iSPNs (left). Quantification of spine density in proximal (n.s. p>0.5) and distal dendrites (n.s. p>0.5) of iSPNs from control and RIM cKO mice (right). Wilcoxon rank-sum test. Summary data are the mean ± SEM.

**Supplemental Figure 12:**
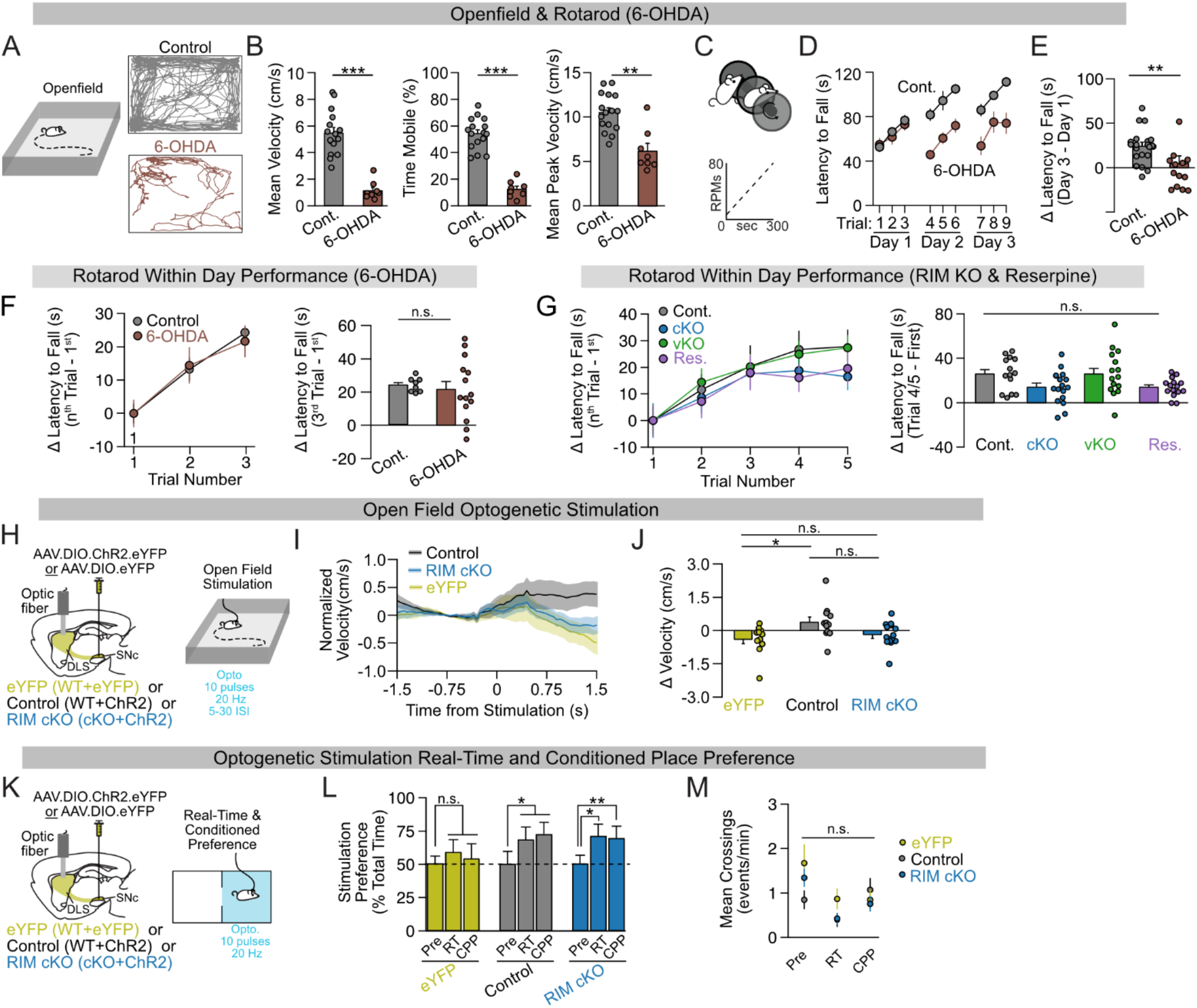
Behavioral controls related to how spatially diffuse and point-to-point dopamine transmission influence motor output and motor learning. A. Schematic of the open field assay and example position plots for a control and 6-OHDA-treated animal. B. Quantification of mean velocity (*** p<0.001), percent time mobile (*** p<0.001), and average peak velocity (** p<0.01) for control and 6-OHDA-treated mice. Wilcoxon rank-sum test. C. Schematic of the accelerating rotarod assay protocol. D. Rotarod performance (latency to fall) over the course of 3 days for control and 6-OHDA-treated mice. E. Quantification of change in performance for control and 6-OHDA-treated mice (** p<0.01). Wilcoxon rank-sum test. F. Summary of the within day changes in performance for control and 6-OHDA-treated mice. Quantification of the changes in performance within days for each group (right, ns p>0.05). Wilcoxon rank-sum test. G. Within day performance changes for control (Cont.), RIM cKO (cKO), RIM vKO (vKO), and reserpine-treated (Res., 0.5 mg/kg) mice (left). Quantification of the within day changes across groups (ns p>0.05). Kruskal-Wallis test (ns p>0.05). H. Schematic of the paradigm for optogenetically stimulating nigrostriatal dopamine axons. Groups included wild type mice expressing eYFP (Control + eYFP) or ChR2 (Control +ChR2) selectively within dopamine neurons and RIM cKO mice expressing ChR2 (RIM cKO + ChR2) within dopamine neurons. I. Change in velocity following light deliver during periods of mobility (>2 cm/s) for the groups described in (H). J. Quantification of the change in velocity following light delivery during periods of mobility (eYFP vs. Control, * p<0.5). Kruskal-Wallis test (* p<0.05) followed by Tukey’s post hoc test (eYFP vs. Control: * p<0.05), eYFP vs. RIM cKO: ns p>0.05, Control vs. RIM cKO: ns p>0.05). K. Experimental paradigm for optogenetic real-time and conditioned place preference. L. Summary of preference for the light-paired chamber at baseline (pre), during real-time place preference (RT), and during conditioned place-preference test session (CPP). (eYFP Pre vs. RT: n.s. p>0.05, eYFP Pre vs. CPP: n.s. p>0.05, Control Pre vs. RT: * p<0.05, Control Pre vs. CPP: * p<0.05, RIM cKO Pre vs. RT: * p<0.05, RIM cKO Pre vs. CPP: **p<0.01). Wilcoxon rank-sum tests. M. Quantification of the average number of crossings per minute across groups for the different test periods (ns p>0.05). Kruskal-Wallis test for each session type. Summary data are the mean ± SEM.

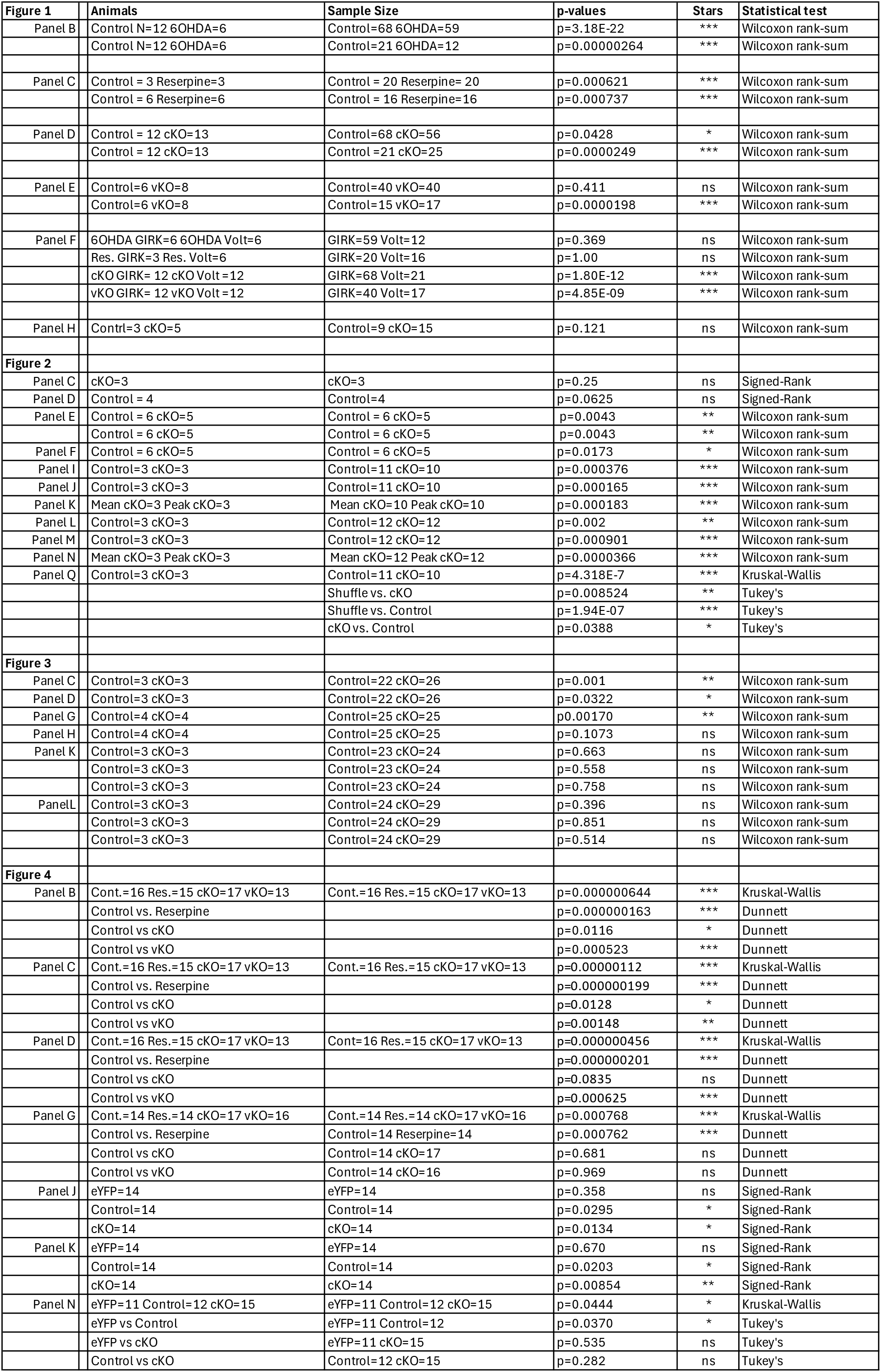
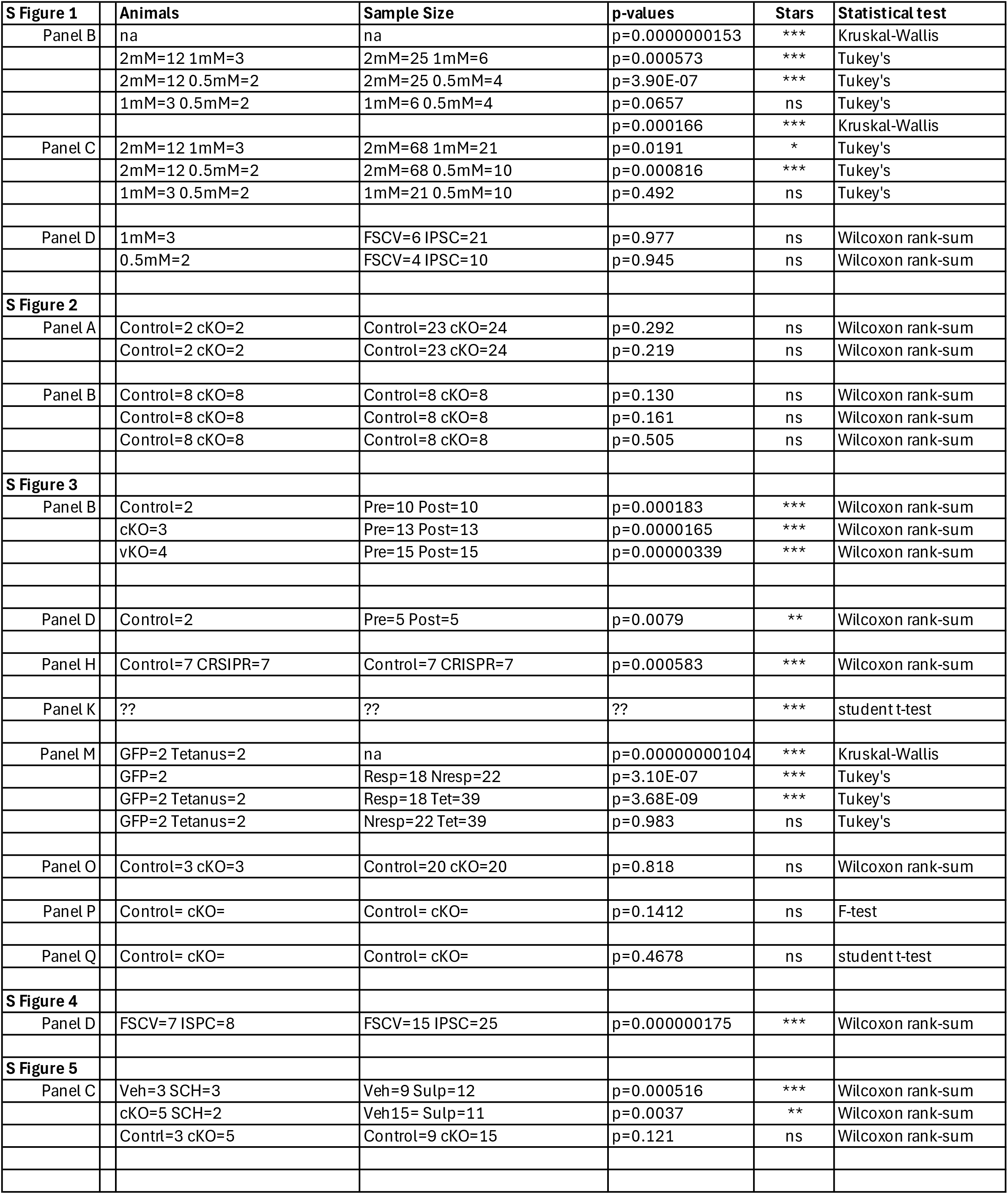
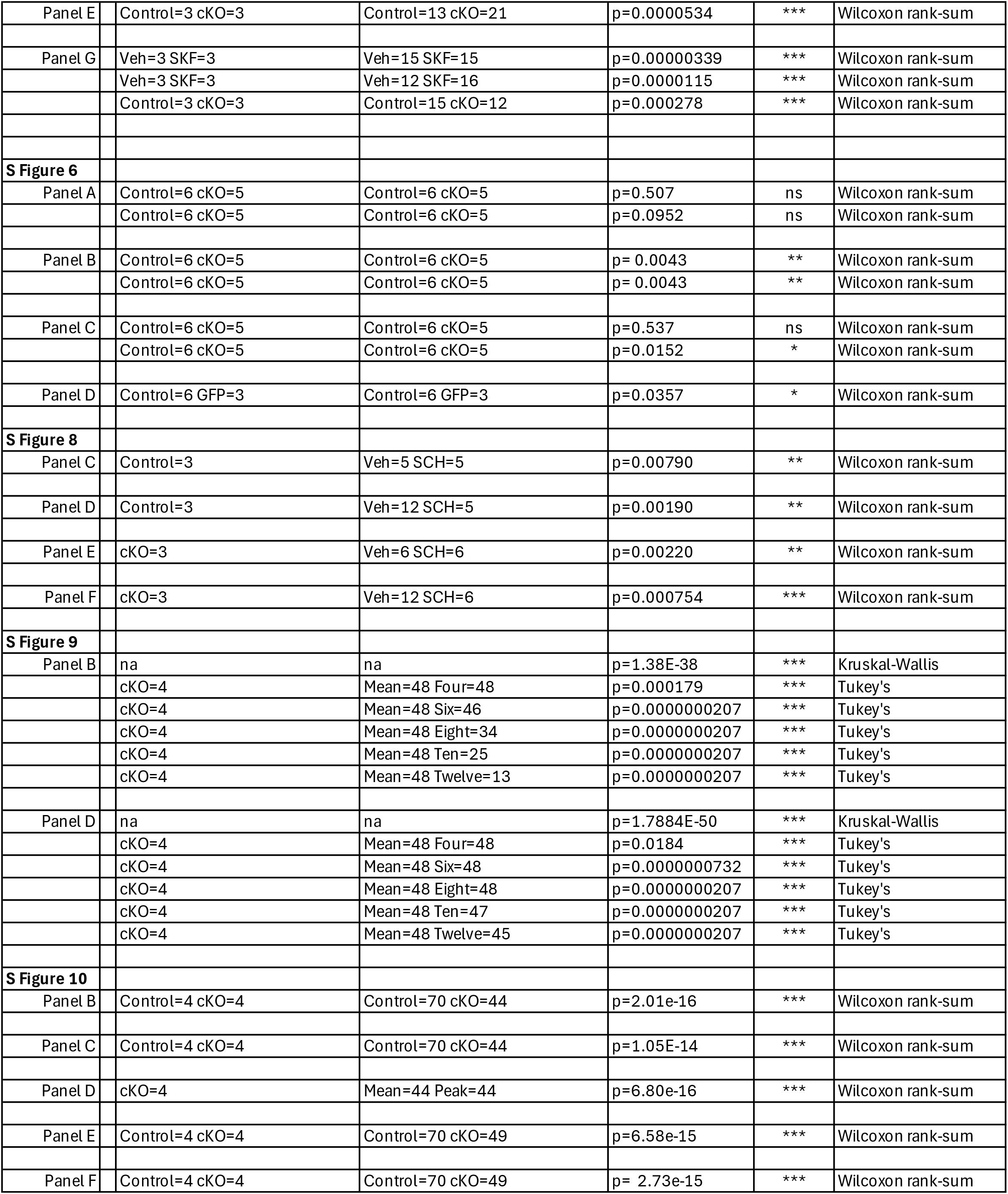
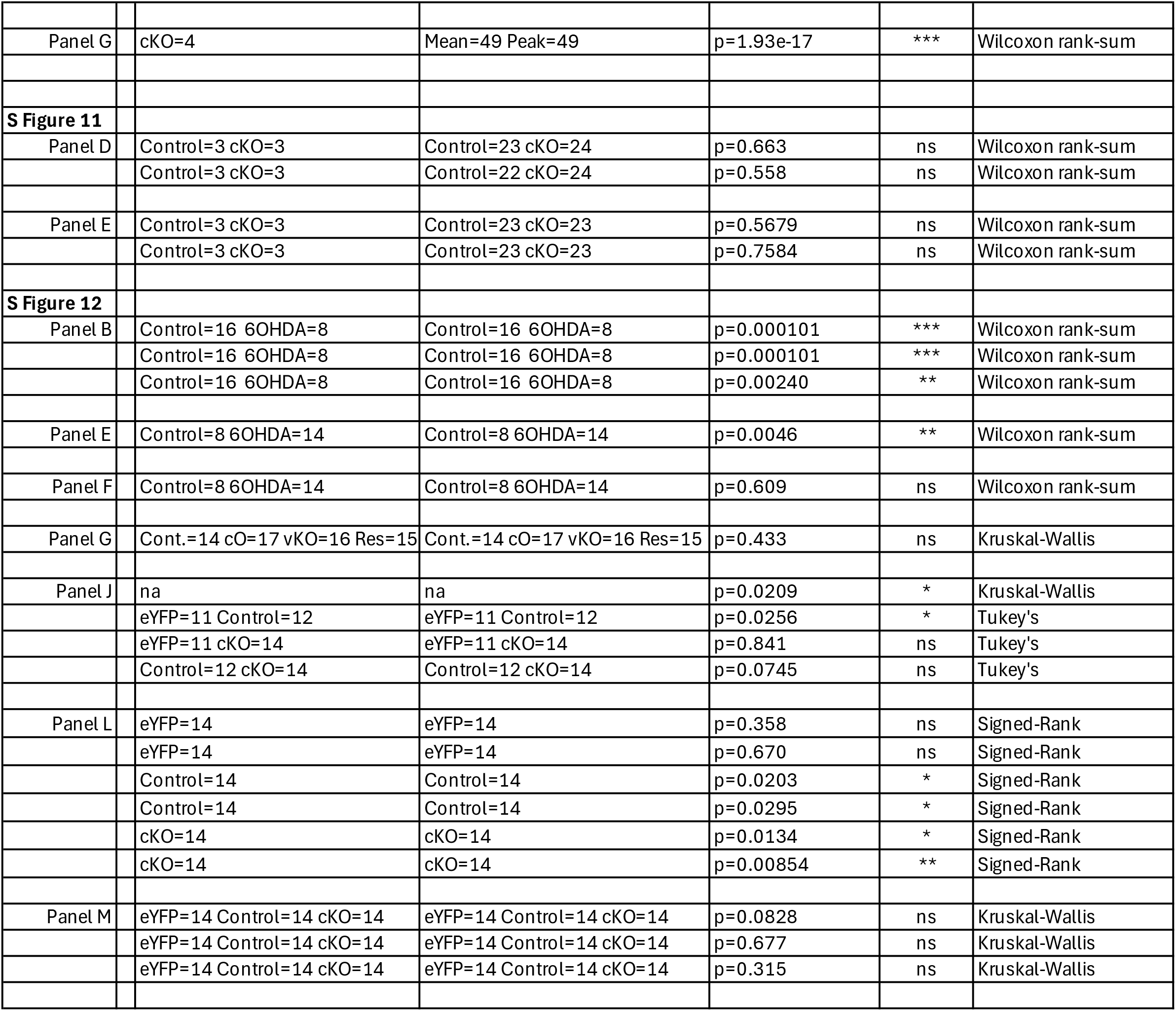

## Notes

### Competing Interest Statement

The authors have declared no competing interest.

